# Transition to the structurally vulnerable nuclear state is an integral part of mouse embryonic development

**DOI:** 10.1101/2023.02.20.529332

**Authors:** Masahito Tanaka, Rin Sakanoue, Atsushi Takasu, Naoko Watanabe, Yuta Shimamoto, Kei Miyamoto

**Affiliations:** Laboratory of Physics and Cell Biology, National Institute of Genetics, Shizuoka 411-8540, Japan; Graduate School of Biology-Oriented Science and Technology, Kindai University, Wakayama, 649-6493, Japan; Department of Genetics, Sokendai University, Shizuoka 411-8540, Japan

## Abstract

Upon fertilization, germ cells are reprogrammed to acquire the ability to develop into an entire organism. Whereas extensive studies have focused on epigenetic reprogramming of chromatin states during development, changes of the nucleus that surrounds chromatin are ill-defined. Here, we show that nuclei become structurally and mechanically vulnerable at the 2-cell stage during mouse embryonic development. The 2-cell stage nuclei are extraordinarily plastic and deformable in contrast to those of 1-cell and 4-cell stages. The mechanically vulnerable nuclear state is attained by autophagy-mediated loss of lamin B1 from the nuclear membrane. This developmentally programmed lamin B1 dynamics is required for chromatin organization and major zygotic genome activation. We thus demonstrate that structural reprogramming of nuclei is a major determinant of embryonic gene expression and acquisition of totipotency.

## Main text

Embryogenesis begins with a pair of haploid genomes that are each packaged in the male and female pronucleus of a zygote. The subsequent cell cleavages allow cell doublings while forming nuclei carrying the diploid genome per blastomere. Within the embryonic nuclei, chromatin states are drastically reprogrammed and this epigenetic reprogramming is key to the transcriptional activation of embryonic genes, referred to as zygotic genome activation (ZGA) (*1, 2*). In mice, major ZGA occurs at the 2-cell stage of embryogenesis and is required for progression of subsequent development. Thus, extensive studies have focused on reprogramming of genomic and chromatin states around this early embryonic stage (*3–8*). In contrast, the state of the nucleus — the “container” of the genome — during this reprogramming process remains a mystery.

The nuclear structure is profoundly affected by nucleoskeletal proteins, including lamins, which assemble underneath the nuclear membrane (*9–11*). Lamins are known to confer mechanical rigidity to the nucleus (*12, 13*), and impairments of lamins can result in abnormally shaped nuclei (*14–16*). Lamins also interact with chromatin and mark transcriptionally inactive loci referred to as lamina-associated domains (LADs) (*11, 17*). The formation of LADs dynamically changes during preimplantation development (*18*) and may thus control ZGA. Heretofore, altered nuclear structure and mechanics have been largely linked to diseases and aging (*19, 20*). A few studies using fly and zebrafish suggested implications of dynamic nuclear structure and mechanics in embryonic development. In early *Drosophila* embryos, nuclear stiffening, mediated by factors other than lamins, parallels transcriptional activation (*21, 22*). A recent work in zebrafish highlighted the maturity of the nuclear pore complex as a determinant of ZGA (*23*). Whether and how lamins control nuclear states during early embryonic development of mammals is not known.

### Nuclei of mouse embryos become plastic and deformable at the 2-cell stage

To investigate the possible changes of nuclear state during early embryonic development, we first analyzed the morphological dynamics of nuclei in living mouse embryos. Using confocal scanning and 3-dimensional (3-D) rendering with histone H2B-mCherry, we found that nuclei at the 1-cell and 4-cell stages were extremely round and spherical (Fig. 1A). On the other hand, nuclei of 2-cell stage embryos exhibited substantial deformation (Fig. 1A and fig. S1A) (see Fig. 1B for quantification). The deformation gradually arose at the early 2-cell stage (20–24 hours post insemination [hpi]) and then became most prominent at the mid 2-cell stage (24–28 hpi) before the smooth, rounded shape was regained towards the end of the 2-cell stage (Fig. 1D and fig. S1, B–E). A micron-sized, concave indentation on one side of the nucleus was typical. The dynamics of nuclear deformation was relatively slow, i.e., the altered shape persisted over several hours with subtle local fluctuations (fig. S1F). In contrast, 1-cell and 4-cell stage nuclei maintained predominant sphericity throughout the cell cycle (Fig. 1, C and E).

**Fig. 1.**
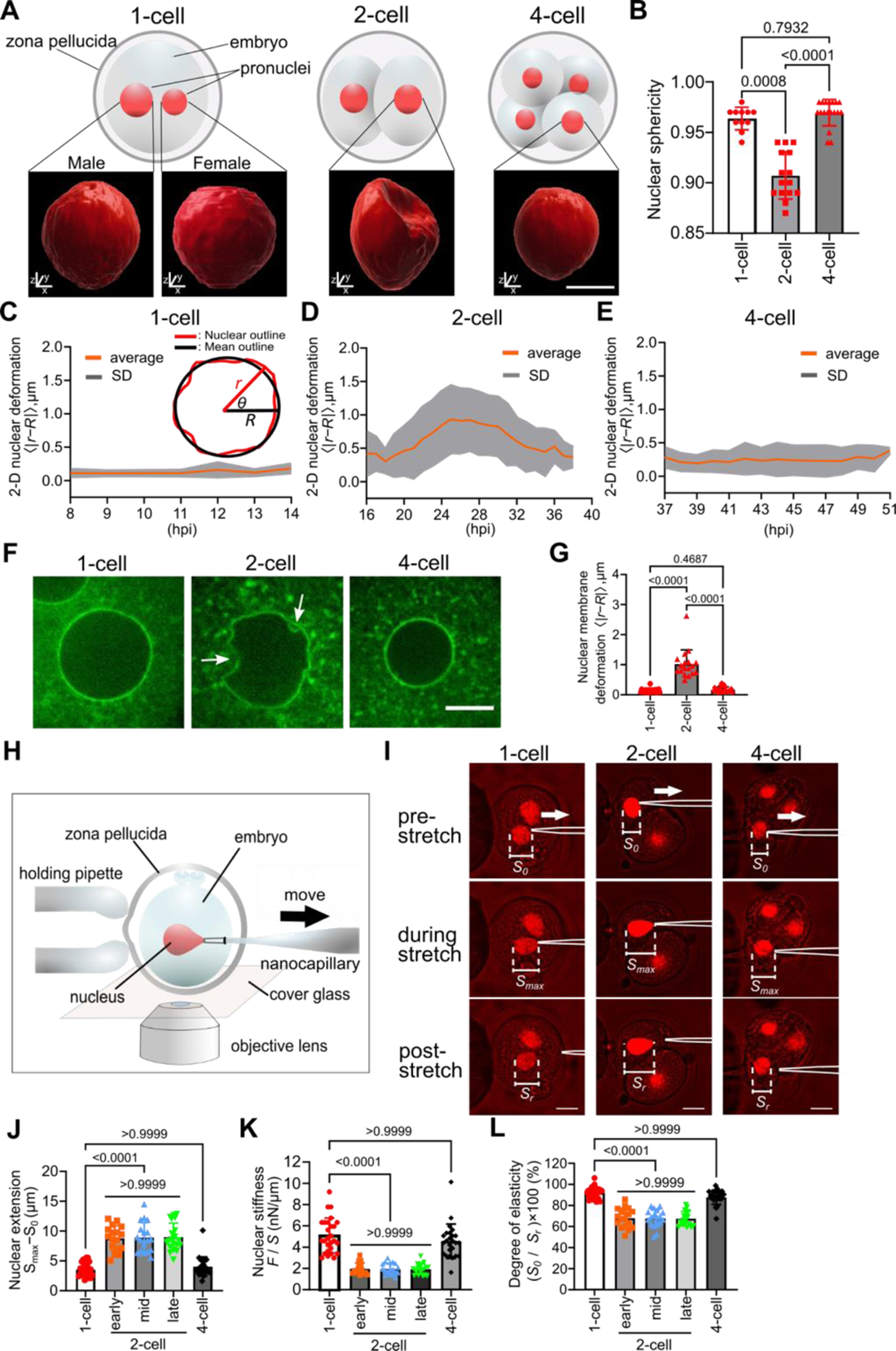
The nucleus undergoes a mechanically deformable state at the 2-cell stage of mouse embryogenic development. (**A**) Schematic (upper) and representative 3-D rendering images (lower; H2B-mCherry) of nuclei at the early cleavage stages. (**B**) Nuclear sphericity. (**C** to **E**) Time courses of nuclear deformation, as analyzed based on displacement of the nuclear periphery (*r*) from the circumference of a perfect circle (radius: *R*) (inset in **C**), at 1-cell (**C**), 2-cell (**D**), and 4-cell stages (**E**). Mean time courses of *n* = 9 embryos (orange) are shown with SD (shaded area). (**F, G**) Nuclear membrane morphology visualized with emerin-EGFP at each embryonic stage (**F**) and the quantified deformation (**G**) (*n* = 21, 18 and 22 nuclei). (**H** to **L**) Nanocapillary-based nuclear stiffness measurement. (**H**) Assay scheme. A single embryo was held with a holding pipette while a nanocapillary was inserted into the embryo to capture the nuclear surface by a weak suction force. The nucleus was then stretched by moving the nanocapillary (black arrow). (**I**) Representative snapshots of 1-, 2-, and 4-cell stage nuclei (H2B-mCherry) stretched by nanocapillaries. The maximal nuclear extension (**J**), nuclear stiffness (**K**) and the degree of elasticity (**L**) were characterized at each embryonic stage by analyzing the nuclear deformation response (as indicated by *S*_0_, *S*_max_ and *S*_r_ in **I**) (colored plots; *n* = 27, 17, 18, 20 and 25 nuclei). Bars are mean ± SD. P values were derived from one-way ANOVA and the Kruskal-Wallis test. All scale bars are 10 μm.

The highly deformed nuclei of 2-cell embryos are suggestive of a loss of structural integrity of nuclear membranes (*24*). We thus visualized the nuclear membrane structure using emerin-EGFP, an inner nuclear membrane marker, or DiI, a lipophilic membrane stain. Our quantitative analysis revealed substantial deformation of the nuclear membrane at the 2-cell stage (Fig. 1, F and G, and fig. S1G), being consistent with the H2B-mCherry imaging. Besides, the surface of the nuclear membrane, which was mostly smooth at the 1-cell and 4-cell stages, was highly disorganized at the 2-cell stage (white arrows, Fig. 1F). At the cytoplasmic-side of the nuclear surface, we observed an accumulation of CHMP7, an ESCRT complex subunit and a marker of membrane damage (fig. S1, H and I) (*25*). Together, these results suggest that the nuclei undergo a drastic structural change at the 2-cell stage during embryonic development.

The disorganized nuclear membrane structure led us to hypothesize that the mechanical stability of the nucleus is also altered in 2-cell embryos. To test this, we measured material properties of embryonic nuclei by developing a nanocapillary-based micromanipulation setup (Fig. 1H). Briefly, the outer surface of the nuclear membrane was captured by a weak suction pressure applied to the nanocapillary (2.1×10^4^ Pa); the nuclear surface was then stretched by an axial translocation of the nanocapillary (Fig. 1I, from top to bottom). Associated with this movement, the nuclei underwent an extensional deformation that gradually increased before being detached from the nanocapillary tip (Fig. 1I, top and middle panels; fig. S2A for typical time course). We then asked to what extent the nuclei could be stretched at each developmental stage. Our quantitative analysis revealed that 2-cell embryo nuclei were >2-times more deformable than those of 1-cell and 4-cell stages (Fig. 1J and fig. S2, B and C). As the propensity of nuclear deformation was not significantly affected by the surrounding cytoplasm (fig. S3, A–F), we could estimate the effective nuclear stiffness based on the extent of maximal nuclear deformation and the amount of capture force (see *Supplementary Materials and Methods*). We found that the nuclear stiffness was lowest at the 2-cell stage (2.0 ± 0.5 nN/µm, *n* = 55 nuclei; pooled sum of early, mid, and late stages) compared to that at the 1-cell and 4-cell stages (5.1 ± 1.6 nN/µm and 4.6 ± 1.6 nN/µm, *n* = 27 and 25 nuclei, respectively) (Fig. 1K). Whereas male pronuclei were slightly more deformable than female pronuclei (4.4 ± 1.4 nN/µm and 5.8 ± 1.7 nN/µm, *n* = 14 and 13 nuclei, respectively), their differences to the 2-cell nuclei were more prominent. The soft and deformable nature of 2-cell stage nuclei persisted throughout the cell cycle while the amount of DNA doubled (early: 2.0 ± 0.5 nN/µm, mid: 2.0 ± 0.5 nN/µm, late: 1.9 ± 0.5 nN/µm; *n*=17, 18, and 20 nuclei, respectively) (Fig. 1, J and K). This 2-cell stage-specific softening of nuclei was further confirmed by employing an alternative assay, which measured the nuclear tension that could withstand a gradually increasing aspiration pressure applied using a wide-bore capillary (*26, 27*) in living embryos (fig. S4). We also noticed that 2-cell stage nuclei had substantial mechanical plasticity; the induced deformation persisted over many minutes after the nuclei were detached from the nanocapillary (bottom middle panel, Fig. 1I). This was in stark contrast to the predominant elasticity of 1-cell and 4-cel stage nuclei, which exhibited rapid shape recovery after the detachment (bottom left and right panels, Fig. 1I; Fig. 1L for quantification). Thus, the nucleus becomes transiently soft and plastic at the 2-cell stage of early embryonic development.

### Autophagy promotes a transient loss of lamin B1 from the nuclear membrane at the 2-cell stage

We explored the molecular mechanism that confers the distinct mechanical and structural properties of the 2-cell embryonic nucleus. A plausible candidate was lamin proteins as they assemble underneath the nuclear membrane. In mouse, there are three major types of lamins, lamin A/C, B1 and B2 (*11, 28, 29*). We thus quantitatively examined the dynamics of each lamin by immunofluorescence. DAPI was used as a DNA counterstain. We found that the membrane localization of lamin A/C showed a gradual decrease as development progressed (fig. S5, A and B), in line with previous work (*28*). The decrease of lamin B2 was also gradual (fig. S5, C and D). In contrast, lamin B1 exhibited a sharp drop at the early-to-mid 2-cell stage and then maintained at a low level, before being raised at the 4-cell stage (Fig. 2, A and B). Immunoblotting revealed a significant reduction of lamin B1 in late versus early 2-cell embryos (Fig. 2C), suggesting a transient degradation of lamin B1 during the 2-cell stage.

**Fig. 2.**
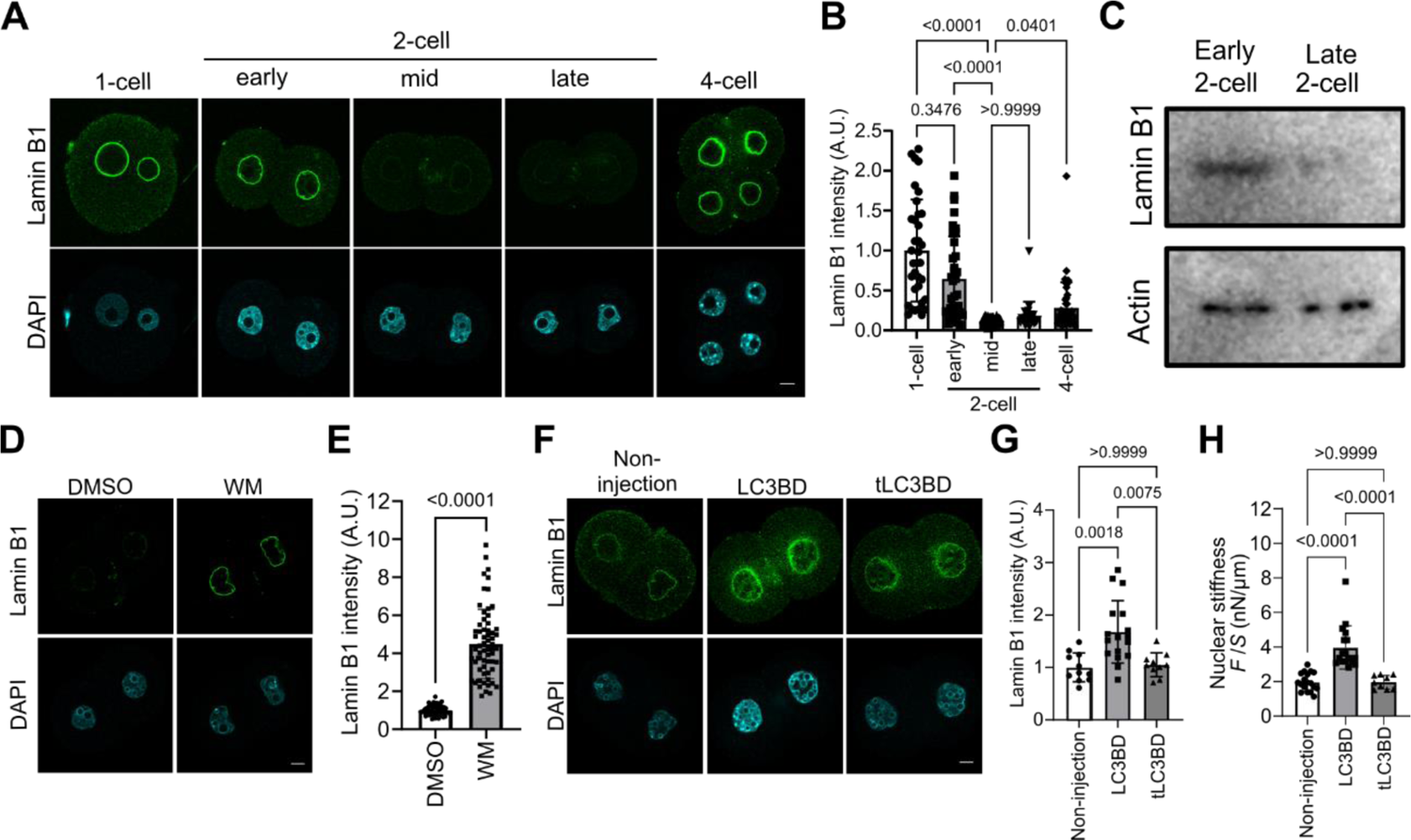
Autophagy-mediated loss of lamin B1 from the nuclear membrane lowers the structural and mechanical stability of embryonic nuclei at the 2-cell stage. (**A, B**) Changes in lamin B1 localization at the nuclear membrane between 1- and 4-cell stages of mouse embryos. (**A**) Representative immunofluorescence images of lamin B1 (upper panels) and DNA (lower panels) at each stage. (**B**) Quantification of lamin B1 signals at the nuclear membrane. Values were scaled to the mean at the 1-cell stage (*n* = 35, 36, 32, 25, and 42 nuclei). (**C**) Immunoblot of early and late 2-cell embryo lysates for lamin B1 and actin (loading control). (**D** to **G**) Effects of autophagy inhibition on lamin B1 nuclear localization. (**D, E**) Effects of wortmannin. (**D**) Representative immunofluorescence images of lamin B1 and DAPI in 2-cell embryos treated with wortmannin (WM), as compared to vehicle control. (**E**) Lamin B1 signals at the nuclear membrane were quantified as in (**B**) (*n* = 56 and 64 nuclei, P <0.001 by Mann–Whitney *U* test). (**F, G**) Effects of LC3 inhibition. (**F**) Representative immunofluorescence images of lamin B1 and DAPI in 2-cell embryos with non-injection control and those injected with mRNA harboring the LC3 binding domain of lamin B1 (LC3BD) or its truncation mutant lacking LC3 binding (tLC3BD) (*n* = 11, 17, 10 nuclei). (**G**) Lamin B1 signal quantification at the nuclear membrane. (**H**) Nuclear stiffness measured using nanocapillaries in embryos with control (*n* = 18; reproduced from Fig. 1K), LC3BD (*n* = 17), or tLC3BD (*n* = 9). Bars are mean ± SD. P values were derived from one-way ANOVA and the Kruskal-Wallis test for Fig.2, B, G, and H.

To determine the molecular mechanism underlying the loss of lamin B1 in 2-cell stage nuclei, we first tested the ubiquitin proteasome pathway which is required for embryonic progression (*30*). However, treatment with MG132, a proteasome inhibitor, did not retain lamin B1 on the nuclear membrane (fig. S6, A and B). We then asked whether autophagy-dependent pathway plays a role in the loss of lamin B1 (*31*). We first used wortmannin, an autophagy inhibitor (*32*). Addition of wortmannin to the culture medium significantly suppressed the 2-cell stage-specific loss of lamin B1 from the nuclear membrane (Fig. 2, D and E). Next, to further verify the contribution of autophagy, we sought to block the interaction of lamin B1 with LC3, a factor responsible for early autophagosome formation (*32*), by injection of mRNA harboring the LC3-binding domain of lamin B1 (hereafter, LC3BD) (*33*). The exogenously expressed LC3BD uniformly distributed across the nuclear interior with no preferential membrane localization (fig. S7A). The loss of lamin B1 in 2-cell stage nuclei was significantly suppressed by LC3BD (Fig. 2, F and G) while nuclear growth (fig. S7B) and other LC3-dependent processes (fig. S7C) were maintained. LC3BD harboring a truncation mutation (tLC3BD), which lacks binding to lamin B1 (*33*), had essentially no effect on the lamin B1 dynamics (Fig. 2, F and G). Importantly, upon inhibition of lamin B1 clearance from the nuclear membrane, 2-cell stage nuclei became significantly more spherical (fig. S8, A and B), with increased stiffness (4.0 ± 1.3 nN/µm) and elasticity (87.6 ± 7.7 % of shape recovery; versus 67.8 ± 8.3 % for non-injection control) (*n* = 17 nuclei, Fig. 2H and fig. S8, C and D). Thus, the loss of structural and mechanical integrity of 2-cell embryonic nuclei is caused by autophagy-mediated clearance of lamin B1 from the nuclear membrane.

### The transient loss of lamin B1 is necessary for major ZGA and embryonic development

We then asked whether the transition to the distinct nuclear state at the 2-cell stage of embryogenesis has any physiological function. Since mouse 2-cell embryos undergo major ZGA (*1*) and the lamin contributes to transcriptional regulation (*34*), we sought to restore nuclear integrity at the 2-cell stage and examine its effect on global gene expression. To this end, 2-cell embryos injected either with the LC3-binding domain of lamin B1 (LC3BD) or the truncated LC3-binding domain (tLC3BD), along with non-injected (Non) embryos were collected at 28 hpi and subjected to RNA-seq analyses (Fig. 3A). In all samples, sequence reads were reproducibly and reliably mapped to the mouse genome (Table S1), but the number of expressed genes (FPKM > 1) significantly decreased in the LC3BD samples (Fig. 3B and Table S1, p < 0.05). We then compared transcriptomes among the different treatments. LC3BD embryos showed lower correlation with other control embryos (fig. S9A, red box). Hierarchical clustering and principal component analyses (PCA) suggested that LC3BD embryos showed distinct transcriptomes from other samples (Fig. 3C and fig. S9B). These results indicate that forced retention of lamin B1 on the nuclear membrane in 2-cell embryos affects global gene expression. We next identified differentially expressed genes (DEGs; padj < 0.05, more than 2-fold difference). Thousands of genes were downregulated in LC3BD embryos when compared to tLC3BD and Non (4680 and 6674 genes, respectively), while very few DEGs were found between tLC3BD and Non (Fig. 3D). Importantly, these downregulated genes well overlapped with major ZGA and 2-cell-activated genes, rather than with minor ZGA and 1-cell-activated genes (*35*) (Fig. 3E, 1623/6674 [24.3%] vs 248/6674 [3.7%], p < 0.01 [Fisher’s exact test]). We then compared the downregulated genes in LC3BD embryos with the recently defined ZGA gene list (*36*) and found that 74.8% of the ZGA genes were repressed in LC3BD embryos (Fig. 3, F and G). Furthermore, Ingenuity Pathway Analysis (IPA) suggested that the downregulated genes in LC3BD embryos showed the mis-regulated canonical pathways (fig. S9C), which highly resembled those reported in major ZGA-deficient embryos (*37*). These results demonstrate that the autophagy-mediated loss of lamin B1 is pivotal for major ZGA.

**Fig. 3.**
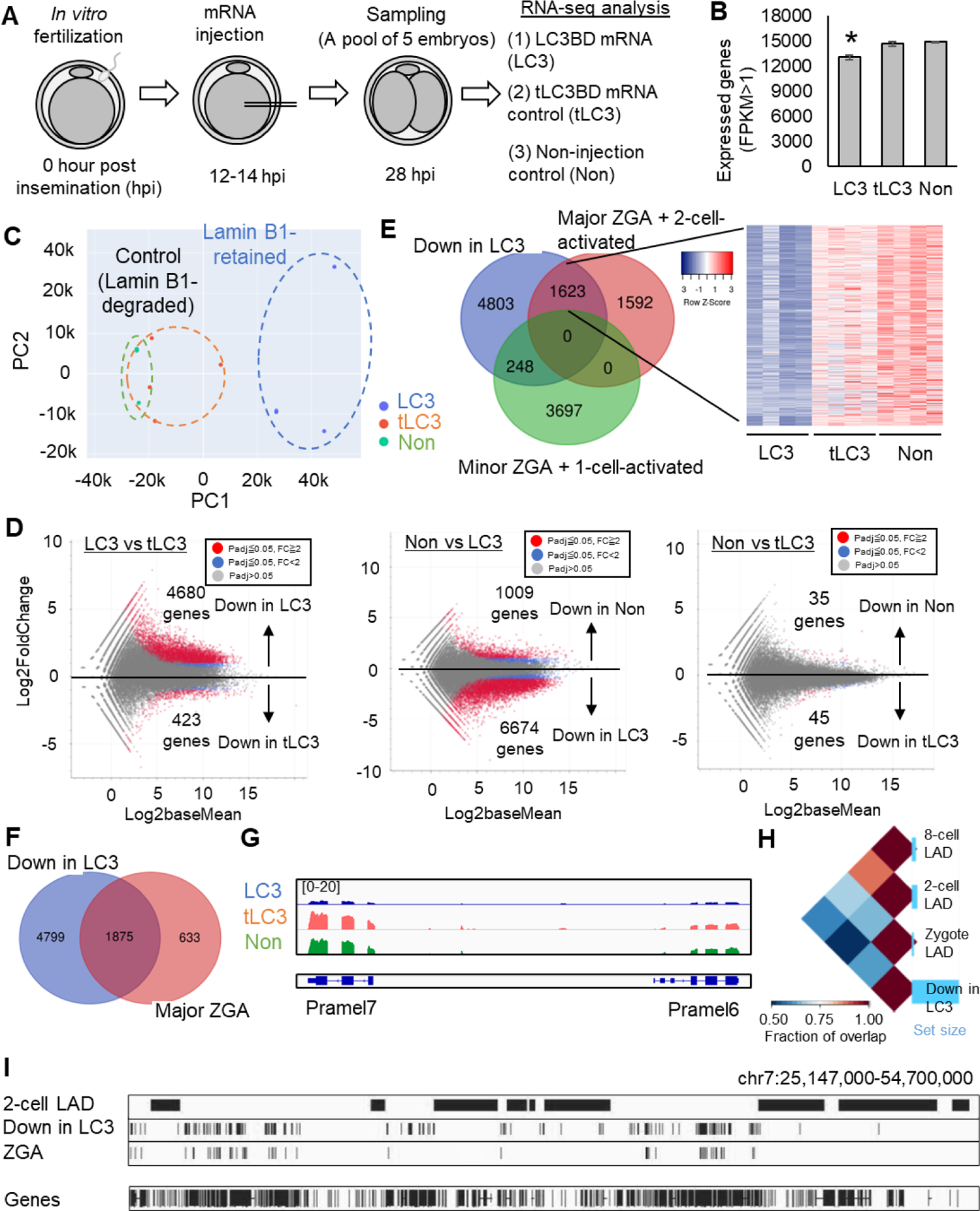
The transient loss of lamin B1 from 2-cell stage nuclei is key to major zygotic genome activation. (**A**) A schematic diagram of RNA-seq analysis of embryos with or without inhibition of lamin B1 loss. (**B**) The number of expressed genes in embryos with different treatments (FPKM: Fragments per kilobase of exon per million mapped reads >1). Mean ± SE are shown. *p < 0.05 (Tukey HSD test). (**C**) PCA of the gene expression profiles. *n* = 12. Blue dots: LC3BD samples (LC3), orange dots: tLC3BD control (tLC3), and green dots: Non-injection control (Non). (**D**) MA plots showing the differentially expressed genes (DEGs). Three different sets of comparison were carried out: LC3BD vs tLC3BD, Non vs LC3BD, and Non vs tLC3BD. The number of downregulated genes (padj < 0.05, fold change > 2) are shown. (**E, F**) Venn diagrams showing the overlap of major ZGA + 2-cell-activated genes (*35*), minor ZGA + 1-cell-activated genes (*35*), and downregulated genes in LC3BD embryos (Non vs LC3) (**E**), and that of major ZGA genes (*36*) and downregulated genes in LC3BD embryos (**F**). A heatmap depicting gene expression levels of overlapping 1623 genes (**E**). (**G**) Track images of RNA-seq reads around Pramel6/7 genes. (**H**) Pairwise intersections of genomic regions among preimplantation LADs (*18*) and downregulated genes in LC3BD embryos. (**I**) Genome browser snapshots depicting genomic locations of 2-cell LADs, downregulated genes in LC3BD embryos, and major ZGA genes.

We further characterized downregulated genes in LC3BD embryos. Genomic Regions Enrichment of Annotations Tool (GREAT) suggested that developmentally essential genes were enriched in the downregulated genes (fig. S10A). The downregulated genes in LC3BD embryos were more correlated with H3K4me3 signals around transcription stat sites (TSSs) than H3K9me3 signals in wild-type 2-cell embryos (fig. S10B), suggesting that these genes should be normally associated with active histone marks for expression at major ZGA. We then examined the relationship between the LC3BD-downregulated genes and LADs in preimplantation embryos (*18*). More than half of the downregulated genes were not associated with the previously identified preimplantation LADs (fig. S10C). Indeed, many of the LC3BD-downregulated genes were not located at the 2-cell LAD (Fig. 3, H and I). Interestingly, most of the major ZGA genes were included in the LC3BD-downregulated genes and those were often excluded from the 2-cell LAD (Fig. 3I). Only 10.8% of major ZGA genes and 13.5% of LC3BD-downregulated genes were associated with 2-cell LADs, while 24.3% of all expressed genes were with 2-cell LADs (fig. S10D). These results suggest that genes expressed in major ZGA are often located outside the LADs, but are largely repressed when lamin B1 retains its localization. Together, the retention of lamin B1 in the 2-cell stage nuclei may lead to abnormal formation of LADs that represses developmentally important major ZGA gene loci.

RNA-seq analyses prompted us to test whether the loss of lamin B1 is related to the transcriptionally permissive chromatin state in 2-cell embryos (*38, 39*). The expression of LC3BD, but not tLC3BD, globally inhibited Pol-II dependent transcription in 2-cell embryos (Fig. 4, A and B). Interestingly, the retention of lamin B1 at the nuclear membrane by LC3BD affected global chromatin organization; DAPI-dense regions were accumulated at the nuclear periphery, while inter-nuclear chromatin became less dense (Fig. 4, C and D). These results suggest that an abnormal accumulation of lamin B1 at the 2-cell stage nuclear membrane biases chromatin towards the transcriptionally repressive state.

**Fig. 4.**
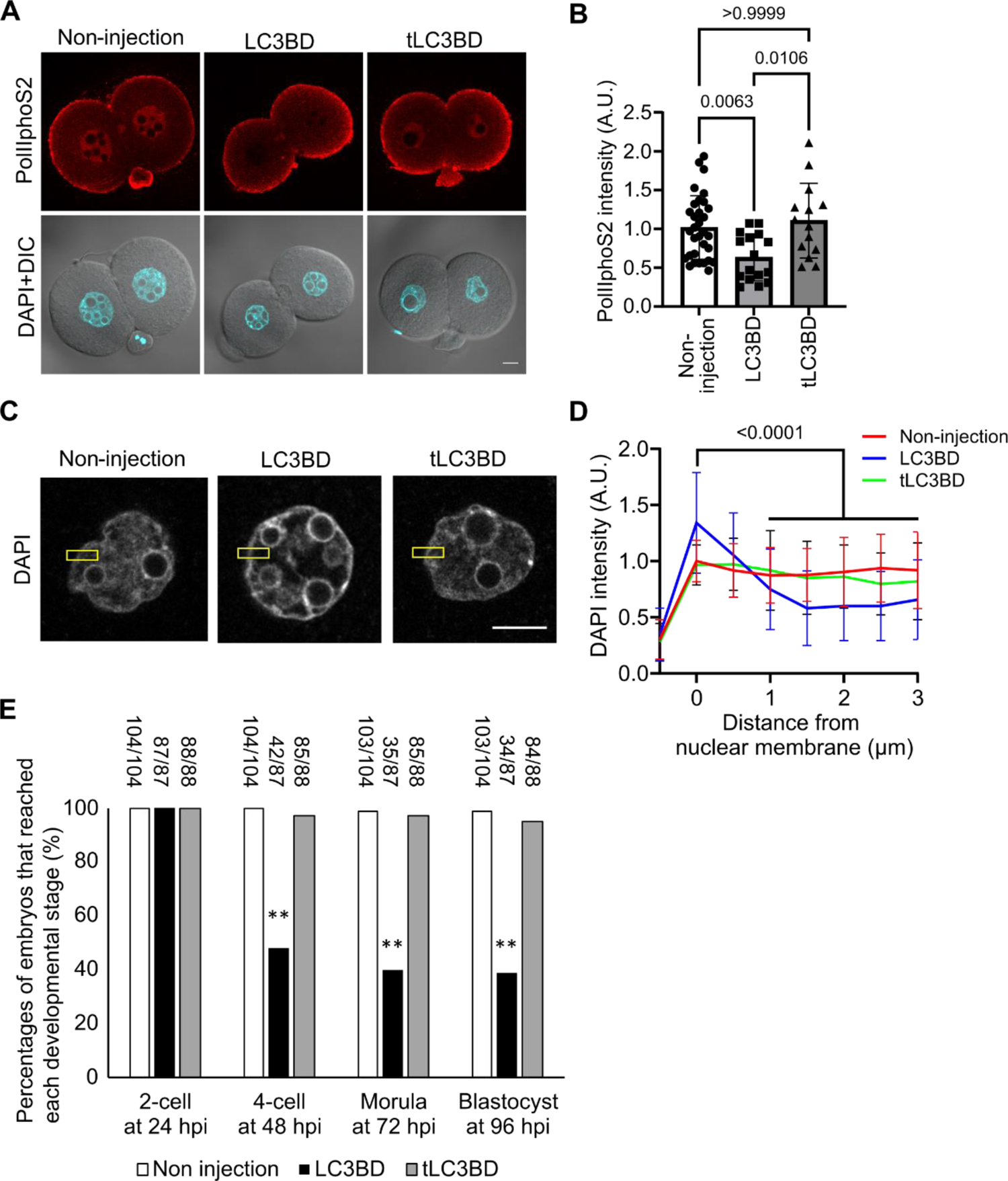
The loss of lamin B1 is required for proper transcriptional activity, chromatin organization and embryonic development. (**A, B**) Transcriptional activity. (**A**) Representative immunostaining images of RNA polymerase II phosphorylated at serine 2 (PolIIphoS2) (upper) and merged images of DAPI and bright-field (lower) in mouse 2-cell embryos at 28 hpi. (**B**) Quantification of the PolIIphoS2 signal within 2-cell embryonic nuclei. *n* = 32 (non-injection), 16 (LC3BD), and 14 (tLC3BD). P values were derived from one-way ANOVA and the Kruskal-Wallis test. (**C, D**) Chromatin organization. (**C**) Confocal images of DAPI-stained nuclei were analyzed for chromatin localization at the vicinity of the nuclear membrane. (**D**) Mean line-scan profiles generated across the nuclear membrane (yellow highlighted areas in **C**). *n* = 56 (non-injection), 44 (LC3BD), and 28 (tLC3BD). P values were derived from one-way ANOVA and the Kruskal-Wallis test. (**E**) Preimplantation development. Embryos of non-injected control and those injected with LC3BD or tLC3BD were cultured until the blastocyst stage. The numbers of embryos that reached each developmental stage are indicated above each bar. P values were derived from the chi-square test (**p < 0.01). All scale bars are 10 µm.

Finally, we asked a developmental role of the distinct nuclear state by experimentally altering it and examining subsequent embryonic development. The mating of heterozygous mutant mice at the lamin B1 locus results in dead pups (*40*) and therefore it has been challenging to investigate lamin B1’s function during early embryonic development. Our approach using LC3BD expression enabled to interfere with endogenous lamin B1 dynamics in early embryos (Fig. 2, F and G). Inhibition of lamin B1 clearance by LC3BD significantly inhibited preimplantation development to the blastocyst stage, whereas tLC3BD did not affect it (Fig. 4E). Thus, the forced restoration of structural and mechanical stability of the 2-cell stage nucleus is detrimental for embryonic development.

## Discussion

We find that the nucleus undergoes dynamic structural reorganization associated with clearance of lamin B1 from the nuclear membrane at the 2-cell stage of mouse preimplantation development. We also demonstrate that this nuclear reorganization alters the nuclear membrane mechanics and is required for proper chromatin assembly and major zygotic genome activation. Previous studies have extensively characterized the epigenetic reprogramming of chromatin states during early embryonic development (*1–8*). Our study emphasizes that the “physical reprogramming” of the nucleus is an integral part of early embryonic development, and suggests that the nuclear structure-oriented regulation can be a driving force for controlling chromatin states in mammals.

Intriguingly, our data reveal that the embryonic nucleus transiently dismantles lamin B1 and lowers mechanical resistance at the 2-cell stage of development. Such a unique physical state of the nucleus is generally considered hazardous for cells, as mechanically fragile nuclei are linked to human diseases such as laminopathies (*41*). However, during embryogenesis, the vulnerable nuclear state is paradoxically required for developmentally essential ZGA. We propose that physical reprogramming of embryonic nuclei leads to the permissive chromatin state for the exceptional transcriptional burst within a short period of time, even though embryos are transiently exposed to a risky state. Mouse early embryos are known to experience potentially deleterious cellular events at the 2-cell stage, such as activation of endogenous retroviruses (*42*) and attenuated DNA damage response (*43*). Future studies will explore how the zygote endures these peculiarities and orchestrates subcellular events for the acquisition of totipotency.

## Acknowledgements

We thank Prof. Kazuo Yamagata, Prof. Yumiko Saga, the members of their laboratories, and Mr. Miyagawa for technical support, Prof. Matsumoto for allowing us to use IPA software, and DNAFORM (Yokohama, Japan) for RNA-seq analyses. We thank Ms. N Backes Kamimura and Mr. James Horvat for proofreading. **Funding:** This work was supported by JSPS KAKENHI Grant Numbers JP19H05751 to K.M. and Y.S., JP22K18362 to Y.S. and 22K15109 to M.T., by The Naito Foundation to K.M., by Takeda Science Foundation to K.M. and Y.S., and by Kindai University Research Grant (19-II-1) and NIG-JOINT (6B2022) to K.M. M.T. is the recipient of a JSPS postdoctoral fellowship. **Author contributions:** Conceptualization: K.M. and Y.S.; Supervision: Y.S. and K.M.; Funding acquisition: K.M. and Y.S.; Methodology: M.T., Y.S., K.M.; Data curation: M.T., R.S., N.W., K.M.; Formal analysis: M.T., R.S., A.T., Y.S., K.M.; Validation: M.T., R.S., Y.S., K.M.; Investigation: M.T., R.S., Y.S., K.M.; Visualization: M.T., R.S., Y.S., K.M.; Writing: M.T., Y.S., K.M. **Author contributions:** The authors declare no competing financial interests. **Data and materials availability:** Requests for plasmids generated in this study should be directed to the corresponding authors. All RNA-seq data have been deposited to the Gene Expression Omnibus (GEO) under accession no. GSE222656. LADs data of mouse zygote, 2-cell and 8-cell embryos were downloaded from GEO accession no. GSE112551. ChIP-seq data of mouse 2-cell embryos were downloaded from GEO accession nos. H3K4me3: GSE72784 and H3K9me3: GSE97778.

## Supplementary Materials

### Materials and Methods

#### Animals

All animal experiments were carried out in strict accordance with the guidelines of Institutional Care and Use of Laboratory Animals. Experimental protocols were approved by the Committee on the Ethics of Animal experiments of Kindai University (Permission#: KABT-31-003) and National Institute of Genetics (Permission#: R4-9). Mice (ICR and B6C3F1/Jc1 [BCF1]) at 5–15 weeks of age (*44*) were purchased from CLEA Japan (Tokyo, Japan), Japan SLC (Shizuoka, Japan), or Kiwa Laboratory Animals (Wakayama, Japan) and maintained in light-controlled, air-conditioned rooms in each mouse facility and were used for *in vitro* fertilization. All mice were sacrificed by cervical dislocation and all efforts were made to minimize suffering and to reduce the number of animals used.

#### *In vitro* fertilization

For obtaining oocytes, female mice were superovulated by intraperitoneal injections of 7.5 IU of pregnant mare serum gonadotropin (PMSG; ASKA Animal Health), followed by 7.5 IU of human chorionic gonadotropin (hCG; ASKA Animal Health) at 48-h post PMSG injection. After the female mice were euthanized at 15-16 h post hCG injection, oocytes arrested at the second meiotic metaphase were collected from the oviducts by an abdominal operation and transferred to a pre-heated medium of Human Tubal Fluid (HTF; ARK Resource). Sperm was collected from the caudal epididymis of male mice for use with *in vitro* fertilization. For fertilization, freshly prepared sperm or cryopreserved and thawed sperm was incubated in HTF medium for 1–1.5 h at 37°C and 5% CO_2_ to allow for capacitation. Subsequently, the sperm suspension was mixed with the medium containing the oocytes, and fertilized eggs were collected 1–2 h after insemination. The cumulus cells attached to the oocytes were removed with 0.1% hyaluronidase at the time of the collection. The embryos were then transferred to drops of preheated KSOM medium (ARK Resource) and cultured at 37°C in air containing 5% CO_2_ until required.

#### Inhibitor treatment

Pharmacological inhibition experiments were performed by treating embryos with the following inhibitors in KSOM medium at the indicated concentrations: Wortmannin (19545-26-7, AdipoGen) (10 µM as the final concentration); MG132 (C2211, Sigma-Aldrich) (5 µM as the final concentration).

#### Antibodies

For immunoblotting assays of p62 and lamin B1, the following antibodies were used: anti-p62 rabbit IgG (PM045MS, MBL), anti-β actin mouse mAb (AC-15, Santa Cruz), anti-GFP chicken IgY (ab13970, abcam), HRP-conjugated goat anti-rabbit IgG antibody (32460, Thermo Fisher Scientific), HRP-conjugated rabbit anti-mouse IgG antibody (61-6520, Thermo Fisher Scientific), HRP-conjugated goat anti-chicken IgY-HRP (A16054, Thermo Fisher Scientific) and anti-lamin B1 mouse IgG (sc-374015, Santa Cruz). For immunostaining assays of lamins and RNA polymerase II, the following antibodies were used: anti-lamin A/C mouse IgG2a (AB_2793218, Active Motif), anti-lamin B1 mouse IgG (sc-374015, Santa Cruz), anti-lamin B2 rabbit IgG (ab151735, abcam), anti-RNA polymerase II phosphorylated at serine 2 mouse IgM (cloneH5: MMS-129R, Covance), goat anti-Mouse IgG (H+L) Cross-Adsorbed Secondary Antibody, Alexa Fluor 488 (A11001, Thermo Fisher Scientific), Goat anti-Rabbit IgG (H+L) Cross-Adsorbed Secondary Antibody, Alexa Fluor 488 (A11008, Thermo Fisher Scientific), Alexa Fluor 594-labeled donkey anti-rabbit IgG antibody (A21207, Thermo Fisher Scientific), and Goat anti-Mouse IgM (Heavy chain) Cross-Adsorbed Secondary Antibody, Alexa Fluor™ 594 (A21044, Thermo Fisher Scientific).

#### Plasmids

pcDNA-emerin-EGFP-polyA was a gift from Yi Zhang (Addgene plasmid #59747) (*45*). pcDNA3.1-Histone H2B-mCherry-polyA and pcDNA3.1-EGFP-polyA were gifts from Dr. Kazuo Yamagata (Kindai University) (*46*). For construction of pcDNA3.1-EGFP-CHMP7-polyA, pcDNA3.1-LC3BD-polyA and pcDNA3.1-EGFP-LC3BD-polyA carrying the LC3-binding domain of mouse lamin B1 (371–459 aa), the corresponding genetic sequences were amplified by PCR from mouse ovary cDNA, which was prepared using SuperScript™ VILO™ Master Mix (11756050, Thermo Fisher Scientific). The amplified fragment was then inserted into the downstream of EGFP of a pcDNA3.1-EGFP vector or replaced with the EGFP sequence of a pcDNA3.1-EGFP vector using In-Fusion HD Cloning Kit (Clontech). The following primer pairs were used for the cloning: chmp7_fp1 (5’–GGGATC CACCGGTCGCCACCATGTGGTCCCCGGAG–3’), chmp7_rp1 (5’–CCCATAGTCGAGCGGCCGCTTAGAGAGTTGGCTCCAATTG–3’), lc3_fp1 (5’–GGGATCCACCGGTCGCCACCATGCTGGACATGGAGATCAG–3’), lc3_rp1 (5’–CATAGTCGAGCGGCCGCTCAGTTCTTCAAGCGAATAAACTTCCCATC–3’), lc3_fp2 (5’–TGGTGGAATTCGCCACCATGCTGGACATGGAGATCAGC–3’) and lc3_rp2 (5’–CATAGTCGAGCGGCCGCTCAGTTCTTCAAGCGAATAAACTTCCCATC–3’). The plasmids carrying truncations (390–438 aa; pcDNA3.1-tLC3BD-polyA) in the LC3-binding domain were constructed from pcDNA3.1-LC3BD-polyA using in-fusion cloning with the following primer pairs: lc3_fp3 (5’–TGGTGGAATTCGCCACCATGCTCTCTCCAAGCCCTTCT TCCC–3’) and lc3_rp3 (5’–GCCCATAGTCGAGCGGCCGCTCAGGCTGAGGCAGAGTGGGAA–3’). All constructed plasmids were verified by DNA sequencing (Eurofins Genetics).

#### mRNA injection

After insemination, fertilized oocytes were collected at 12–14 hpi for mRNA injection. Cumulus-free embryos were transferred to drops of M2 medium (ARK Resource) for injection. mRNAs were injected using a piezo manipulator (Prime Tech, Tsukuba, Japan). The final concentrations of injected mRNA are as follows; 500 ng/μl LC3BD + 25 ng/μl EGFP and 500 ng/μl truncated LC3BD (tLC3BD) + 25 ng/μl EGFP. EGFP mRNA was mixed to confirm successful injection. After injection, embryos were cultured in KSOM medium at 37°C in air containing 5% CO_2_. mRNAs were synthesized from plasmid vectors linearized by restriction enzyme digestion at 37°C overnight and then processed with mMESSAGE mMACHINE T7 Transcription Kit (AM1344, Thermo Fisher Scientific). The synthesized mRNAs were purified using RNeasy Plus Mini Kit (74034, QIAGEN) and subjected to agarose gel electrophoresis and spectrophotometry to check the purity (criteria: OD 260/280 ratio of >2.0) prior to injection into oocytes. The purified mRNAs were flash frozen in liquid nitrogen, stored at –80°C, and used within 6 months.

#### Live cell imaging

Live cell imaging was performed using an inverted fluorescence microscope (Ti-U, Nikon) equipped with a motorized sample stage (MS-2000, Applied Scientific Instruments), a piezo-driven objective scanner (PIFOC, Physics Instruments) with a piezo controller (E-665, Physics Instruments), a stage-top incubator (TIZ, Tokai Hit), a spinning-disk confocal unit (CSU-X1, Yokogawa) with two excitation lasers (488 nm and 561 nm; OBIS LS, Coherent), a silicone-immersion objective lens (UPlanSApo 60× 1.30 NA, Olympus) and a sCMOS camera (Neo4.1, Andor). To begin with, the fertilized embryos prepared as above were injected with mRNA (20 ng/μl of histone H2B-mCherry or 300 ng/μl of emerin-EGFP) at 1–2 h post-insemination and then transferred to drops of KSOM medium (10 µl each), which was supplemented with 0.00025% of polyvinyl alcohol (PVA; P8136-250G, Sigma-Aldrich) (*47*) and placed on a glass-bottom culture dish (P35G-1.5-14-C, MatTek) filled with paraffin oil (26137-85, Nacalai Tesque). The embryos were cultured in a cell culture incubator before being transferred to the microscope’s stage-top incubator, which was maintained at 37°C and filled with humidified air containing 5% CO_2_. Time-lapse imaging was started at 8 h post insemination with 1 h intervals while the optical section of embryos was acquired across individual nuclei (step size: 5 µm; total travel: 60 µm). The exposure of embryos (exposure time: 100 ms each) to the excitation lasers was minimized by shuttering the illumination using image acquisition software (NIS elements v5.0, Nikon). The laser power at the objective’s focus was set to 0.2 mW and 0.3 mW for 488 nm and 561 nm lasers, respectively. Under these conditions, embryos developed with no noticeable photodamage (∼82% reached to the blastocyte stage; versus ∼80% for control samples cultured in an incubator; *n* = 11 and 47 embryos, respectively). Nuclear membrane images (fig. S1G) were taken by injecting DiI (D3911, Thermo Fisher Scientific) into embryos after dissolving the dye in oil (23-0510-5, Sigma Ardrich) and immediately before imaging. For 3-D reconstruction of nuclear shape morphology (Fig. 1A and fig. S1A), confocal slices were acquired at 1 µm steps. The image rendering was performed using Imaris software (v9.8.2, Zeiss).

#### Immunofluorescence imaging

IVF embryos were fixed in 4% PFA/PBS at room temperature for 20 min, and were washed three times by 0.01% PVA/PBS. Samples were next treated with 0.5% triton X-100/PBS at room temperature for 20 min, followed by three times incubation with 3% BSA/PBS each for 10 min at room temperature. The samples were then incubated with primary antibodies (anti-Lamin B1[B-10) mouse IgG [1:100, sc-374015, Santa Cruz], anti-Lamin B2 rabbit IgG [1:100, ab151735, Abcam], anti-Lamin A/C mouse IgG [1:200, 39087, Active Motif], or anti-RNA polymerase II phosphorylated at serine 2 mouse IgM [1:100, clone H5: MMS-129R, Covance] at 4°C overnight. Following three times of washes by 3% BSA/PBS, samples were further incubated in the dark with goat anti-Mouse IgG (H+L) Cross-Adsorbed Secondary Antibody, Alexa Fluor 488 (1:2,000; A11001, Thermo Fisher Scientific), goat anti-Rabbit IgG (H+L) Cross-Adsorbed Secondary Antibody, Alexa Fluor 488 (1:2,000; A11008, Thermo Fisher Scientific), Alexa Fluor 594-labeled donkey anti-rabbit IgG antibody (1:2,000; A21207, Thermo Fisher Scientific), or goat anti-Mouse IgM (Heavy chain) Cross-Adsorbed Secondary Antibody, Alexa Fluor™ 594 (1:2,000; A21044, Thermo Fisher Scientific) at room temperature for 1 h. The samples were washed with 3% BSA/PBS three times and then mounted on slides using VECTASHIELD Mounting Medium containing DAPI. The fluorescence signals were observed using a LSM800 microscope, equipped with a laser module (405/488/561/640 nm) and GaAsP detector, using the same contrast, brightness, and exposure settings within each set of experiments. Z-slice thickness was determined by using the optimal interval function in the ZEN software.

#### Nuclear mechanics measurements

The mechanical properties of nuclei at each embryonic stage (i.e., 1-, 2-, and 4-cell stages) were measured in living embryos using single glass nanocapillaries, which were held by a three-axis, low-drift hydraulic micromanipulator (MHW-3, Narishige) attached to the microscope’s body. The nanocapillaries were prepared by fabricating borosilicate glass tubes (B100-75-10, Sutter Instruments) using a capillary puller (PD-10, Narishige) such that the tip had a nearly uniform taper with an inner diameter of ∼1 μm (1.06 ± 0.02 µm, *n* = 5). To begin with, embryos injected with mRNAs (20 ng/μl of H2B-mCherry or 300 ng/μl of emerin-EGFP) were transferred to drops of M2 medium (ARK Resource), which was placed in a glass-bottom, open-top culture chamber covered with paraffin oil (26137–85, Nacalai Tesque). The chamber was then placed on the microscope’s sample stage and a single embryo was captured by using a large holding pipette (inner diameter: ∼10 µm), which approached from one side of the embryo. A nanocapillary was then inserted from the opposite side of the embryo and its tip was contacted to the nuclear surface such that the nucleus could be captured at its surface by a weak suction pressure applied to the nanocapillary. The level of the suction pressure was controlled using a syringe pump (PHD ULTRA, Harvard apparatus), and a digital manometer (PPA101-M, SMC), which were connected to the distal end of the nanocapillary via a vacuum tube, was used to monitor the pressure level. The pressure level (2.1 × 10^4^ Pa) was such that the nuclear surface could sustain its attachment to the nanocapillary tip but could not be aspirated. Finally, the nucleus was stretched by moving the nanocapillary position away from the embryo’s center (Fig.1I) at a velocity with which the viscous resistance from the cytoplasm could be minimal. At the typical movement speed of 15 µm/min, the viscous resistance on the nucleus as estimated according to the Stokes-Einstein equation (*48*) was 0.05–0.16 nN. This value was >100-times smaller than the capture force generated at the nanocapillary tip (∼17 nN), as estimated from the product of the applied pressure (2.1×10^4^ Pa) and the nanocapillary’s aperture size (∼0.8 µm^2^; = *π* (*d*/2)^2^, where *d* is the nanocapillary’s inner diameter). Associated with the nanocapillary’s movement, the nucleus gradually increased the extent of deformation and then detached from the nanocapillary before reaching the embryo’s edge. The extent of the maximal extension that individual nuclei exhibited was used to determine the degree of nuclear deformability. The measurement was performed with nuclei of 1-, 2-, and 4-cell embryos using approximately the same micromanipulation parameters (i.e., pressure level, nanocapillary size and nanocapillary movement speed). A fraction of embryos (n = 33 of 140) had nuclei that were entirely aspirated off by the applied pressure; this was observed irrespective of the examined cell stages and thus excluded from subsequent analysis.

The mechanical properties of nuclei were also examined using an alternative technique (fig. S4), which was also based on glass capillaries but eliminated of the nuclear translocation within the cytoplasm. This was achieved by using the same micromanipulation setup as above but with the pressure level that was gradually increased instead of being maintained constant; the critical tension above which the nuclear membrane was aspirated inside a glass capillary (inner diameter: 2.09 ± 0.12 µm, *n* = 13) was measured. Specifically, following the physical contact of the capillary tip with the nuclear surface, the suction pressure was raised at a constant rate of 6 × 10^3^ Pa/min. The capillary’s position was fixed throughout this procedure. The nucleus could withstand the applied aspiration pressure and maintain its overall shape until it yielded to the pressure. The pressure level was constantly monitored using a digital manometer (Testo 521, Testo) and the value at which the nuclear surface started to migrate inside the capillary (>1 µm displacement from the tip) was determined by examining the nuclear image. The measurements were performed at 22 ± 1°C and completed within 60 min, over which no noticeable change in the nuclear mechanics was observed. The objective lens of 40× (Plan Apo, 0.95 NA; Nikon) was used for imaging.

#### Western blot

p62 and lamin B1 were immunoblotted on embryonic lysates. For p62 blotting, embryos were first injected with mRNA (500 ng/µl of the EGFP-LC3BD) at 12–14 h post-insemination and then collected at the 2-cell stage (at 32 hpi). Before lysis, the zona pellucida was removed by acidified Tyrode’s (T1788, Sigma-Aldrich) and washed three times in PBS containing 0.01% PVA. Sixty embryos were then soaked with Tris-Glycine SDS sample buffer (LC2676, Novex) containing 2-mercaptoethanol (161-0710, BioRad), and 500 ng BSA (Albumin Standard, 23209, Thermo Fisher Scientific) as a loading control and heated at 100 °C for 5 min. For lamin B1 blotting, 34 IVF embryos were collected at 21 and 32 hpi, and lysed in SDS sample buffer (Wako). The lysis was done by heating at 100 °C for 3 min. Proteins were separated on 10% polyacrylamide gel or e-PAGEL HR (5-20%) gel (HER-T520L, ATTO) and then transferred to PVDF membrane (1704272, BioRad) using Transblot SD (BioRad) or a Trans Blot-Turbo transfer system (1704150J1, BioRad). Following blocking with 3% BSA-containing PBS with 0.2% Tween or SuperBlock Blocking Buffer (37535, Thermo Fisher Scientific) and 0.05% Tween (161-0781, BioRad), the membrane was blotted sequentially with the following primary antibodies: anti-Lamin B1(B-10) mouse IgG (1:1000, sc-374015, Santa Cruz) or anti-p62 rabbit IgG (1:1000, PM045MS, MBL), anti-actin rabbit IgG (1:1000, A2066, Merck), and then anti-GFP Chicken IgY (1:5000, ab13970, abcam) (for p62 blotting only). Each blotting was carried out overnight at 4°C, followed by 1-h staining with the secondary antibody at room temperature; HRP-conjugated goat anti-rabbit IgG antibody (1:3000, 32460, Thermo Fisher Scientific), HRP-conjugated rabbit anti-mouse IgG antibody (1:3000, 61-6520, Thermo Fisher Scientific), and HRP-conjugated goat anti-Chicken IgY-HRP (1:5000, A16054, Thermo Fisher Scientific). The membrane was stripped off using Restore Western Blot Stripping Buffer (21059, Thermo Fisher Scientific) following each blot. For p62 blotting, the membrane was pre-stained with Ponceau S (P7170, Sigma-Aldrich) and then washed with DPBS after image acquisition. Blotting was performed using ECL™ Prime Western Blotting System (Amersham, RPN2232) or SuperSignal West Femto according to the manufacturer’s protocol (34094, Thermo Fisher Scientific). Images were acquired using Gel Doc ZE imager (BioRad).

### Data analysis

#### Measurement of nuclear shape

The nuclear shape was analyzed in 2-D or 3-D using confocal sections of nuclei acquired during live cell imaging. For the analysis of 2-D deformation, the contour of each nucleus, which was visualized with H2B-mCherry or emerin-EGFP, was detected at the equatorial plane by using an Image J plug-in (Jfilament2D; https://www.lehigh.edu/~div206/jfilament/download.html; curve type = contour, energies = Gradient (edges), others are as default) after processing the image with a median filter for noise reduction (2 × 2 pixel). The centroid position of the nucleus (*x*_0_, *y*_0_) was then determined from the detected nuclear contour, and the average radius (*R*) of the nucleus was calculated according to the following equation:

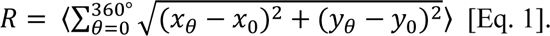

Here, (*x*_θ_, *y*_θ_) are the positions of the nuclear contour at the angle *θ* (0° ≤ θ < 360°) relative to the nuclear center position (*x*_0_, *y*_0_). Subsequently, a perfect circle of the radius *R* was drawn from the nuclear center and the deviation of the nuclear contour from the circle was calculated at each angle *θ* using the LEAP Image J plug-in (https://www.lehigh.edu/~div206/LEAP/index.html). The extent of nuclear deformation (*D*) was determined as a mean deviation of the nuclear contour from the perfect circle according to the following equation:

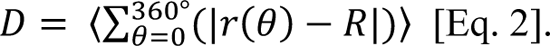

Here, *r* (*θ*) is the distance of the nuclear contour from the nuclear center at the angle *θ*. For analyses of H2B channel images, the nuclear contour was detected at their outer edges. For analyses of emerin channel images, the nuclear contour was detected at their inner edges because their outer edges were largely disorganized.

The 3-D nuclear shape was analyzed using Imaris (v9.8.2, Oxford Instruments). Each stack of confocal slices acquired at 1 µm steps across the nucleus was used to reconstruct the 3-D nuclear morphology and the sphericity was calculated using the Imaris Surfaces function.

#### Measurement of nuclear mechanical properties

The deformability of nuclei (*S*) was determined according to the following equation:

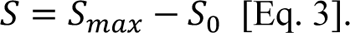

Here, *S*_0_ is the initial resting length of the nucleus measured prior to stretch, and *S*_max_ is the maximal length that the nucleus reached before detachment from the nanocapillary. Each nuclear length was determined along the micromanipulation axis by measuring the distance between the nanocapillary’s contact point and its opposite point across the nucleus. The analysis was performed for multiple embryos imaged with either H2B-mCherry or emerin-EGFP and consistent results were obtained (fig. S2C). The initial nuclear size of the 1, 2, and 4-cell embryos was 20.2 ± 2.7 µm (*n* = 27), 14.8 ± 1.8 µm (*n* = 18), and 15.5 ± 1.6 µm (*n* = 25), respectively.

The nuclear stiffness (*K*) was estimated based on the maximal nuclear deformation (*S*) and the amount of force (*F*) applied at the nanocapillary tip, according to *K* = *F*/*S*. The force *F* (= 17 nN) was determined as the product of the applied suction pressure (2.1×10^4^ Pa) and the area at which the pressure was applied (i.e., the bore size of the nanocapillary) (0.8 ± 0.1 µm^2^, *n* = 5). The bore size was measured for each nanocapillary by filling its interior with a dye solution. The speed of nuclear translocation associated with the nanocapillary movement was 0.26 ± 0.15 µm/s (*n* = 107). The viscous resistance associated with this movement (*F*_v_; 0.05–0.16 nN) was estimated according to the Stokes-Einstein equation, which was given by *F*_v_ = 6πηaV. Here, *η* is the cytoplasmic viscosity (1.2–3.6 Pa-s) (*49, 50*), *a* is the average nuclear radius, and *V* is the average speed of nuclear translocation. The estimated viscous resistance from the cytoplasm was >100 times smaller than the force holding the nucleus by the nanocapillary. The cytoplasmic elasticity had little influence on our analysis as well, as the nuclei did not return to the original position after detachment from the nanocapillary (fig. S3, D to F).

The extent of nuclear elasticity (*R*) (Fig. 1L) was determined based on its deformation recovery, by calculating the ratio of the nuclear length prior to stretch (*S*_0_) to the nuclear length after detaching from the nanocapillary and reaching a new steady-state (*S*_r_), according to the following equation: *R* = *S*_0_⁄*S*_r_ × 100. The new steady nuclear length *S*_r_ was determined by fitting the relaxation time course of the nuclear length to the Exponential Recovery function in Image J.

The mechanical resistance of nuclei, as examined using an increased suction pressure from the ∼2 µm bore capillaries (fig. S4), was determined by simultaneously monitoring the nuclear shape and the pressure level at which the nuclear surface was aspirated inside the capillary (> 1µm from the tip). Time-lapse images of nuclei visualized with H2B-mCherry were used to monitor the nuclear shape.

#### Immunofluorescence image quantification

Equatorial planes manually selected from confocal slices of immunostaining images were used to quantify fluorescence intensities of PolIIphoS2 and lamins in image J. For analysis of PolIIphoS2, mean intensities of PolIIphoS2 signals were calculated within individual 2-cell nuclei and background signals were subtracted from the nuclear signals. For analysis of lamin, the following procedure was taken. First, in each two-color image, which was composed of lamin immunofluorescence and DAPI counterstain, the nuclear contour was detected by an automatic thresholding of the DAPI signal after median filtering (2 × 2 pixel). A band was then drawn with the line width of ± 1 μm from the detected nuclear contour and transferred to the lamin image, in which the integrated fluorescence intensity within the band (*I*_total_) was measured. The background fluorescence (*I*_bg_) was calculated from intensities inside and outside the nucleus (*i*_bg_in_ and *i*_bg_out_, respectively; ± 5 μm area from the detected nuclear contour excluding the band) according to the following equation:

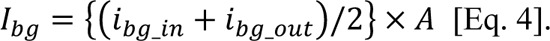

Here, *A* is the total area of the band placed to cover the nuclear contour. Finally, the mean signal intensity of lamin at the nuclear contour (*I_lamin_*) was determined by subtracting the total background (*I*_bg_) from the integrated signal intensity (*I*_total_) measure within the band.

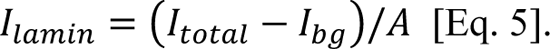

### RNA-sequencing (RNA-seq) analysis

At 28 hpi, a pool of five IVF embryos at the 2-cell stage per condition was treated with acid Tyrode, followed by three times washes with 0.1% BSA/PBS, and were transferred to a 0.2 mL tube containing 9.5 μL of reaction buffer (Z5013N, Takara) to make a final volume of 10.5 μL. After cell lysis in the buffer, 1 μL of the solution was removed and, instead, 1 μL of the appropriately diluted ERCC RNA Spike-In Mix was added (Thermo Fisher Scientific, 4456740). SMART-seq library preparation was performed using SMART-Seq® HT Kit (Z4437N, Takara) and Nextera XT DNA Library Preparation Kit (FC-131-1096, illumine, San Diego, CA) according to the vendor’s instruction. We followed the previously published protocol for RNA-seq of mouse embryos (*51*). The analysis was performed with four biological replicates.

The paired-end sequencing (150 bp + 150 bp) was obtained by the Illumina NextSeq 2000 platform. Raw reads were first subjected to filtering to remove low quality reads using Trimmomatic (*52*). Reads of less than 20 bases and unpaired reads were also removed. Furthermore, adaptor, polyA, polyT and polyG sequences were removed using Trim Galore! (https://www.bioinformatics.babraham.ac.uk/projects/trim_galore/). The sequencing reads were then mapped to the mouse genome (mm10) using STAR (*53*). Reads on annotated genes were counted using featureCounts (*54*). RNA-seq reads were visualized using Integrative Genome Viewer (*55*). FPKM values were calculated from mapped reads by normalizing to total counts and transcript. Differentially expressed genes (DEGs, padj < 0.05, Fold Change > 2) were then identified using DESeq2 (*56*) by normalizing with ERCC RNA Spike-In counts. A hierarchical clustering of the read count values was performed using hclust in TCC (unweighted pair group method with arithmetic mean: UPGMA). Principal-component analysis (PCA) of the global gene expression profile was performed using scikit-learn and the plots were depicted using plotly. Each gene list was further subjected to an Ingenuity Pathway Analysis (IPA; QIAGEN, Redwood City, CA). Using IPA, enriched canonical pathways were investigated. Using GREAT (http://great.stanford.edu/public/html/), functions of DEGs were predicted. The genomic regions that overlapped between the published LADs (GSE112551) and our identified DEGs were analyzed by using BedSect (*57*) or Intervene (*58*). To visualize normalized enrichment of ChIP-seq read counts (H3K4me3: GSE72784 and H3K9me3: GSE97778) (*6, 59*) at LC3-downregulated genes in mouse 2-cell embryos, heatmaps were depicted using computeMatrix and plotHeatmap in deepTools (*60*).

### Statistical analysis

Statistical significance was calculated by the Kruskal-Wallis test for Fig.1, B, G, and J-L, Fig.2, B, G, and H, Fig. 4, B and D, fig. S1, D, E, and I, fig.S3, B and E, fig.S4C, fig.S5, B and D, fig.S6B, fig. S8, B and D, by Mann–Whitney *U* test for Fig.2E, fig. S2C and fig. S7B, by chi-square test for Fig. 4E, by Tukey HSD test for Fig. 3B, and by Fisher’s exact test for Fig. 3E. Mean and standard deviation (or standard error) values are reported in the associated figure captions. Sample size was determined based on our preliminary experiments and no statistical method was used. Embryos were randomly selected for imaging, mechanics and gene expression analyses. Blinding was applied for the analysis of image data. Embryos that showed abnormal appearance such as degenerated embryos were not subjected to analyses.

### Data availability

The RNA-seq data produced in this study are available in the following database: Gene Expression Omnibus (GSE222656).

## Supplementary Figures

**Figure S1.**
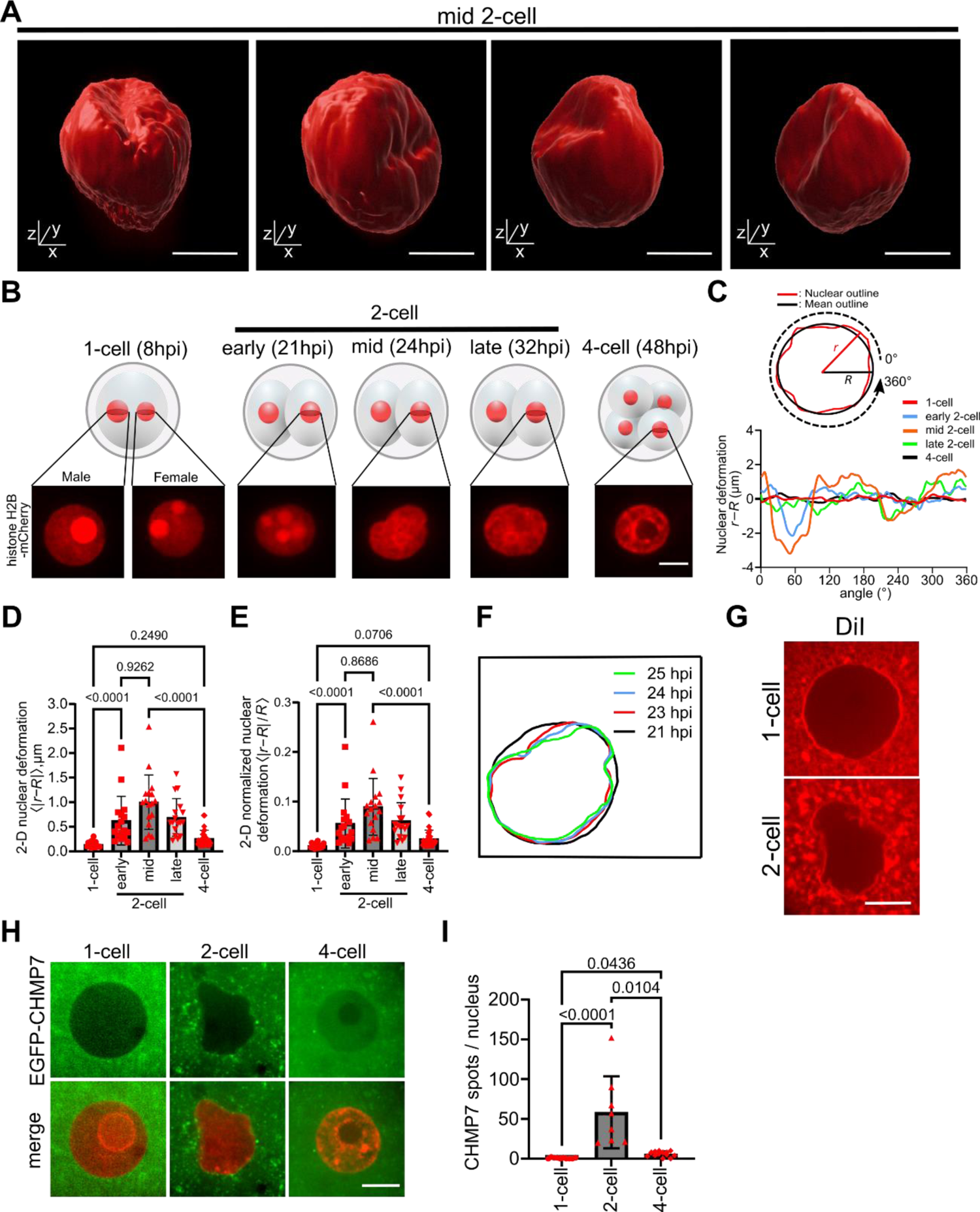
Nuclear deformation at the 2-cell stage of mouse embryonic development. (**A**) Additional examples of the 3-D reconstruction images generated as in Fig. 1A using confocal live cell slices of mouse 2-cell stage nuclei captured at 24 hpi, confirming the consistency of the observed nuclear deformation. Scale bars, 10 µm. (**B**) Confocal snapshots showing time-dependent changes of 2-D nuclear morphology visualized using H2B-mCherry and captured in a developing single embryo. The bright circular structures within 1-cell and early 2-cell nuclei are nucleolus precursor bodies. Scale bar, 10 µm (consistent across the images). (**C**) Line profiles of nuclear deformation over time. The displacement of the nuclear periphery position (*r*) (red solid line) from a perfect circle of radius *R* (= 〈r(θ)〉^360°^, black solid line) was expanded over 360° along the nuclear circumference (upper schematic). The deformation profiles generated at each cell stage from single nuclei are shown (colored lines). (**D, E**) Comparison of nuclear deformation analyzed using 2-D confocal images. The absolute nuclear deformation (**D**) and the relative magnitude of deformation normalized to nuclear size (**E**) at different cell stages are shown. Measurements were performed at the equatorial plane of each nucleus and the mean magnitude of deformation along the nuclear circumference was calculated. The values in (**D**) are 0.14 ± 0.06 µm (1-cell), 0.62 ± 0.50 µm (early 2-cell), 1.00 ± 0.56 µm (mid 2-cell), 0.69 ± 0.38 µm (late 2-cell) and 0.25 ± 0.17 µm (4-cell). The values in (**E**) are 1.1 ± 0.1 × 10^-2^ (1-cell), 5.6 ± 4.9 × 10^-2^ (early 2-cell), 8.9 ± 5.7 × 10^-2^ (mid 2-cell), 6.1 ± 3.7 × 10^-2^ (late 2-cell) and 2.5 ± 1.7 × 10^-2^ (4-cell) (mean ± SD, *n* = 9 embryos). (**F**) Time-dependent change of nuclear shape morphology over 4 h during the mid-to-late 2-cell stage. (**G**) Nuclear membrane structures visualized using DiI at the 1-cell versus 2-cell stages. (**H, I**) ESCRT localization as determined by CHMP7 accumulation. (**H**) Representative confocal snapshots of nuclei at each embryonic stage visualized with H2B-mCherry (red) and EGFP-CHMP7 (green). The images were generated as superposition of five consecutive Z-sections (step size: 1 µm) and the number of the CHMP7 foci (white arrows) at the vicinity of the number membrane was counted (**I**). The values are 0.6 ± 0.7 (1-cell), 58.4 ± 45.4 (2-cell), and 5.9 ± 3.3 foci per nucleus (4-cell) (mean ± SD, *n* = 10, 8 and 14 nuclei). P values were derived from one-way ANOVA and the Kruskal-Wallis test. Scale bar, 10 μm (consistent across the images).

**Figure S2.**
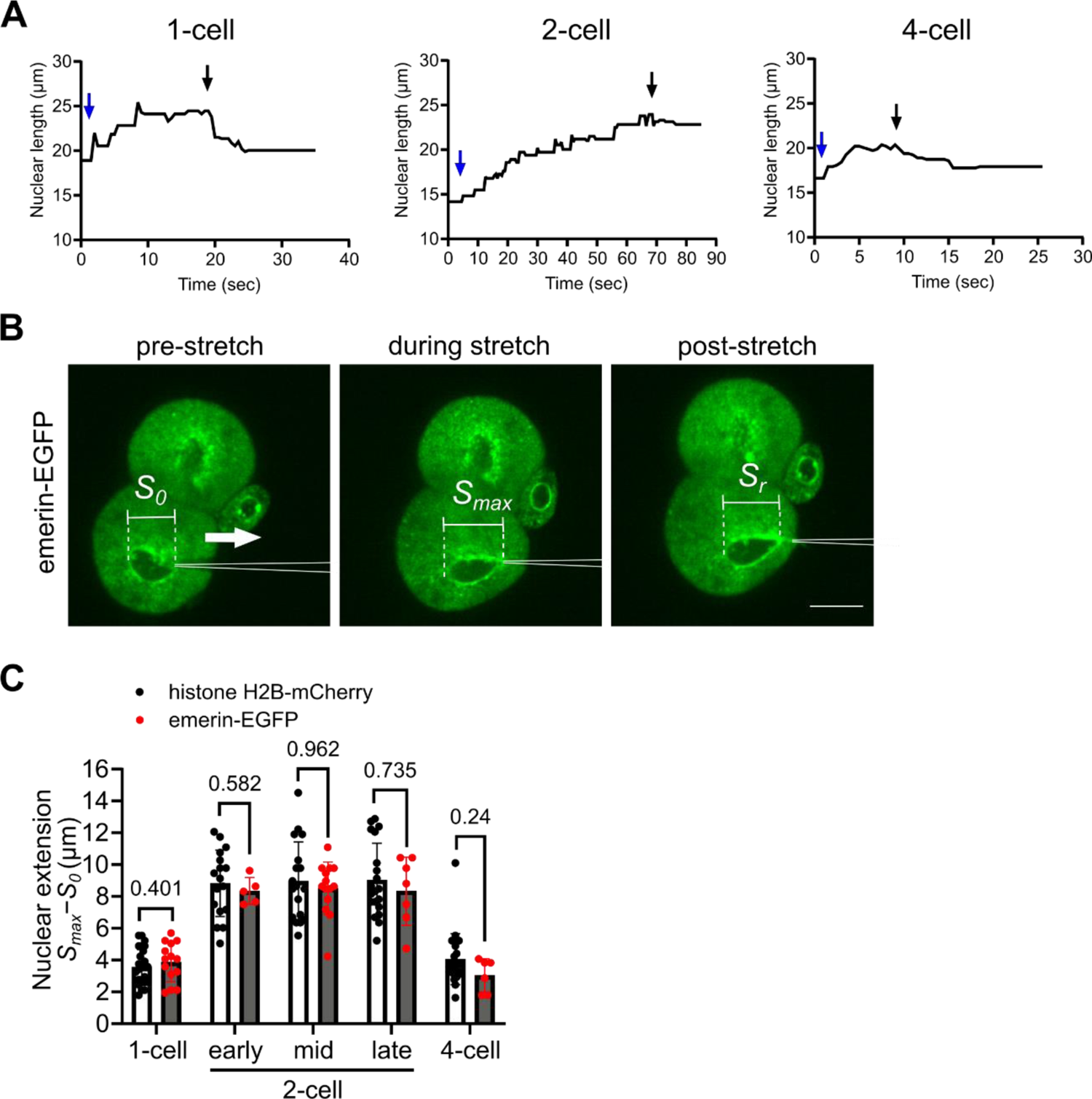
Nanocapillary-based nuclear deformability measurement in early mouse embryos. (**A**) Time courses of nuclear extension (*S*, vertical axis) that developed in response to the stretching micromanipulation performed using nanocapillaries. Representative traces at the indicated cell stages are shown. The downward arrows indicate the time at the onset of nuclear stretch (blue) and the time at which the nucleus detached from the nanocapillary (black). (**B, C**) Comparison of nuclear deformability examined using histone H2B-mCherry and emerin-EGFP. Shown in (**B**) are representative confocal snapshots of a 2-cell stage embryo injected with emerin-EGFP mRNA and subjected to micromanipulation as in Fig. 1H. The nanocapillary position is outlined with white solid line. Scale bar, 20 μm (consistent across the images). (**C**) Quantitative analysis confirmed that the maximal nuclear extension examined at each embryonic stage using emerin-EGFP and H2B-mCherry were indistinguishable from each other (P > 0.05 by Mann–Whitney *U* test). The values determined using emerin-EGFP were 3.9 ± 1.2 µm (1-cell), 8.4 ± 0.9 µm (early 2-cell), 8.5 ± 1.7 µm (mid 2-cell), 8.3 ± 2.1 µm (late 2-cell) and 3.0 ± 1.1 µm (4-cell) from *n* = 14, 5, 14, 7, and 6 nuclei, respectively (mean ± SD). The values determined using H2B-mCherry (replotted from Fig. 1J) were 3.6 ± 1.1 µm (1-cell), 8.8 ± 2.1 µm (early 2-cell), 9.0 ± 2.5 µm (mid 2-cell), 9.0 ± 2.3 µm (late 2-cell) and 4.1 ± 1.6 µm (4-cell) from *n* = 27, 17, 18, 20 and 25 nuclei, respectively.

**Figure S3.**
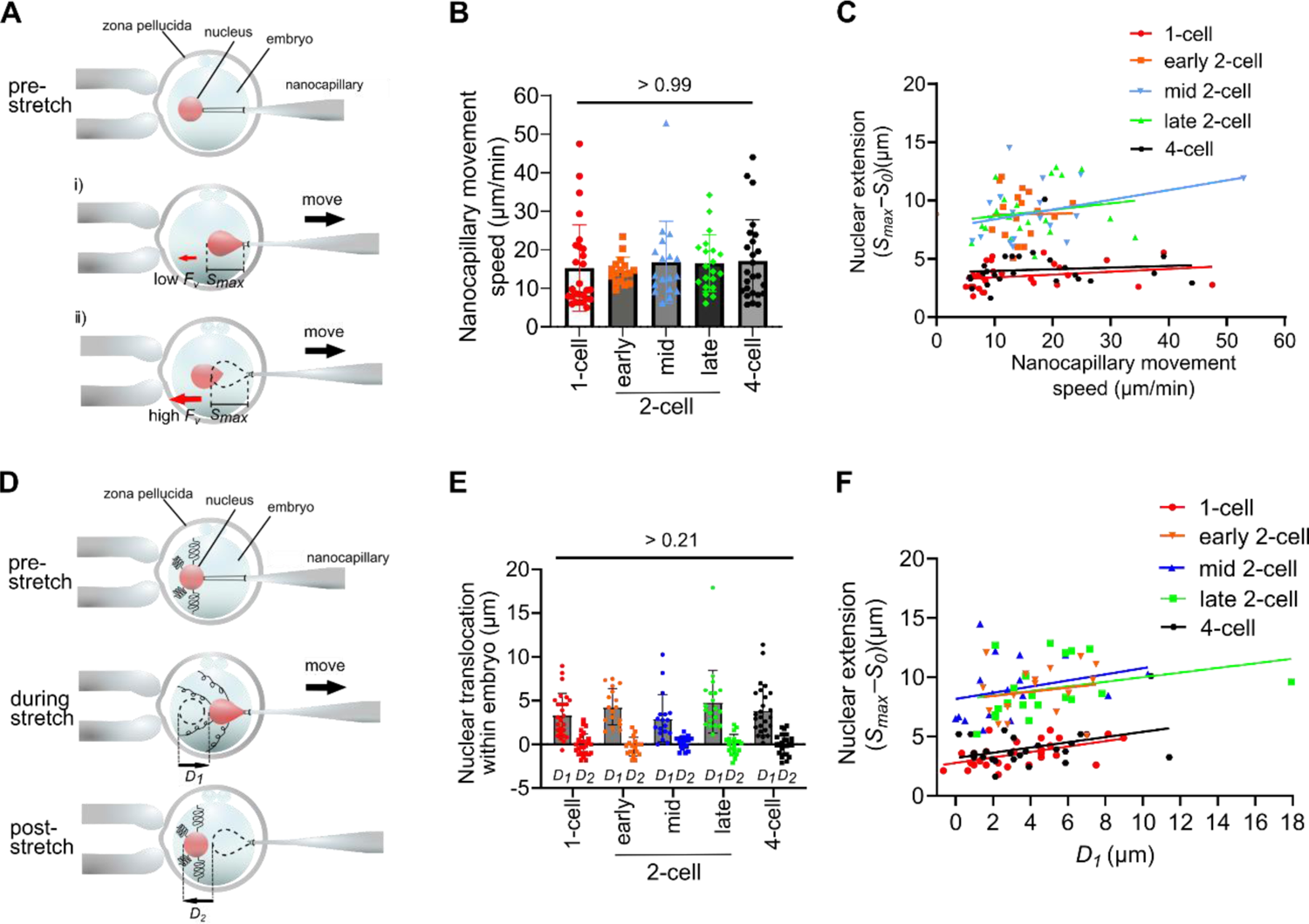
Cytoplasmic mechanical properties do not significantly affect the measured nuclear deformability. (**A** to **C**) Influence of cytoplasmic viscosity on the nanocapillary-based nuclear deformability measurement. As the nucleus was translocated within the cytoplasm, a viscous drag may arise at the vicinity of nuclear surface. This drag force should become larger when the nucleus was pulled at a faster velocity, causing an early detachment of the nucleus from the nanocapillary and potentially influencing the measurement (**A**). The average speed of nanocapillary movement used for the measurement was comparable across different cell stages (P > 0.99 by one-way ANOVA and Kruskal-Wallis test) (**B**). A variation in the speed of nanocapillary movement was recognizable across each measurement (individual plots in **B**), allowing for plotting the observed magnitude of nuclear deformation against the pulling speed for individual trials (**C**). The deformation magnitude was nearly independent of the pulling speed, indicating negligible influence of cytoplasmic viscosity on the measured nuclear deformability. The colored lines in **C** are linear regression slope; 0.02, −0.01, 0.08, 0.06, and 0.01 for 1-cell (red), early 2-cell (orange), mid 2-cell (blue) and late 2-cell (green) and 4-cell (black), respectively. (**B**). Bars are mean ± S.D. Values are 15.3 ± 11.2 µm/min (1-cell), 14.5 ± 3.5 µm/min (early 2-cell), 16.8 ± 10.6 µm/min (mid 2-cell), 16.5 ± 7.4 µm/min (late 2-cell) and 17.1 ± 10.7 µm/min (4-cell) (*n* = 27, 17, 18, 20 and 25 nuclei, respectively). (**D** to **F**) Influence of cytoplasmic elasticity on the nanocapillary-based nuclear deformability measurement. If there is a predominant elastic element within the cytoplasm, the nuclear translocation associated with the nanocapillary movement (*D*_1_) should strain the element and cause a generation of a restoring force on the nucleus, which moves the nucleus back to the initial position in the embryo following detachment from the nanocapillary (*D*_2_) (**D**). Our analysis showed that *D*_2_ was nearly zero in the measurement at any embryonic stages examined, suggesting no predominant elasticity within the cytoplasm (P > 0.21 by one-way ANOVA and Kruskal-Wallis test) (**E**). The measured nuclear deformation was nearly independent of the magnitude of nuclear translocation (*D*_1_) (**F**), further supporting that the cytoplasmic elasticity minimally influenced the measured propensity of nuclear deformation. The colored lines in **F** are linear regression slope; 0.22, 0.18, 0.26, 0.19, and 0.22 for 1-cell (red), early 2-cell (orange), mid 2-cell (blue) and late 2-cell (green) and 4-cell (black), respectively.

**Figure S4.**
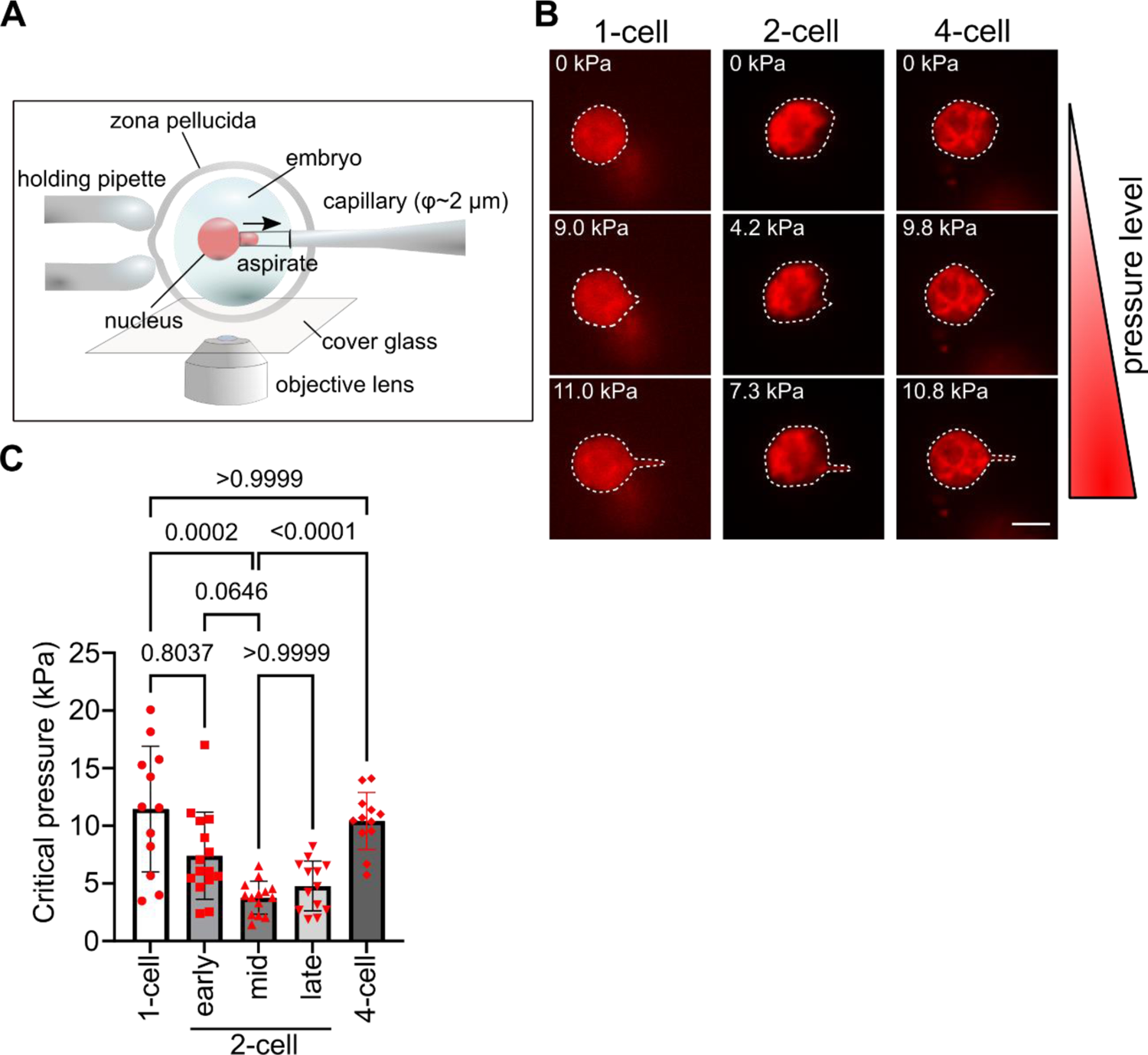
Low mechanical resistance of 2-cell embryonic nuclei against aspiration force. (**A**) A schematic diagram shows an alternative measurement of nuclear mechanics using capillary suction. A capillary (bore size: ∼2 µm) was first contacted to the surface of the embryonic nucleus. A negative pressure was then applied with a constant increment of 6×10^3^ Pa/min and the critical pressure level at which the nuclear membrane was aspirated inside the capillary (>1 µm displacement from the tip) was measured. (**B**) Confocal live-cell imaging snapshots of nuclei at each embryonic stage visualized with histone H2B-mCherry and subjected to an increased suction pressure from the capillary. Each nucleus, whose surface was subjected to the pressure (top panels), showed local deformation around the capillary tip (middle panels) and then started to be aspirated inside it (bottom panels) at different pressure levels (upper left corner of each image). Dotted lines highlight nuclear contours. Scale bar, 10 μm (consistent across the images). (**C**) Quantification of the pressure level required for aspirating the nuclear surface at each embryonic stage. Bars are mean ± S.D. The values are 11.5 ± 5.5 kPa (1-cell), 7.4 ± 3.8 kPa (early 2-cell), 3.8 ± 1.4 kPa (mid 2-cell), 4.8 ± 2.2 kPa (late 2-cell), and 10.4 ± 2.5 kPa (4-cell) (*n* = 12, 15, 14, 13 and 12 nuclei, respectively). P values were derived from one-way ANOVA and the Kruskal-Wallis test.

**Figure S5.**
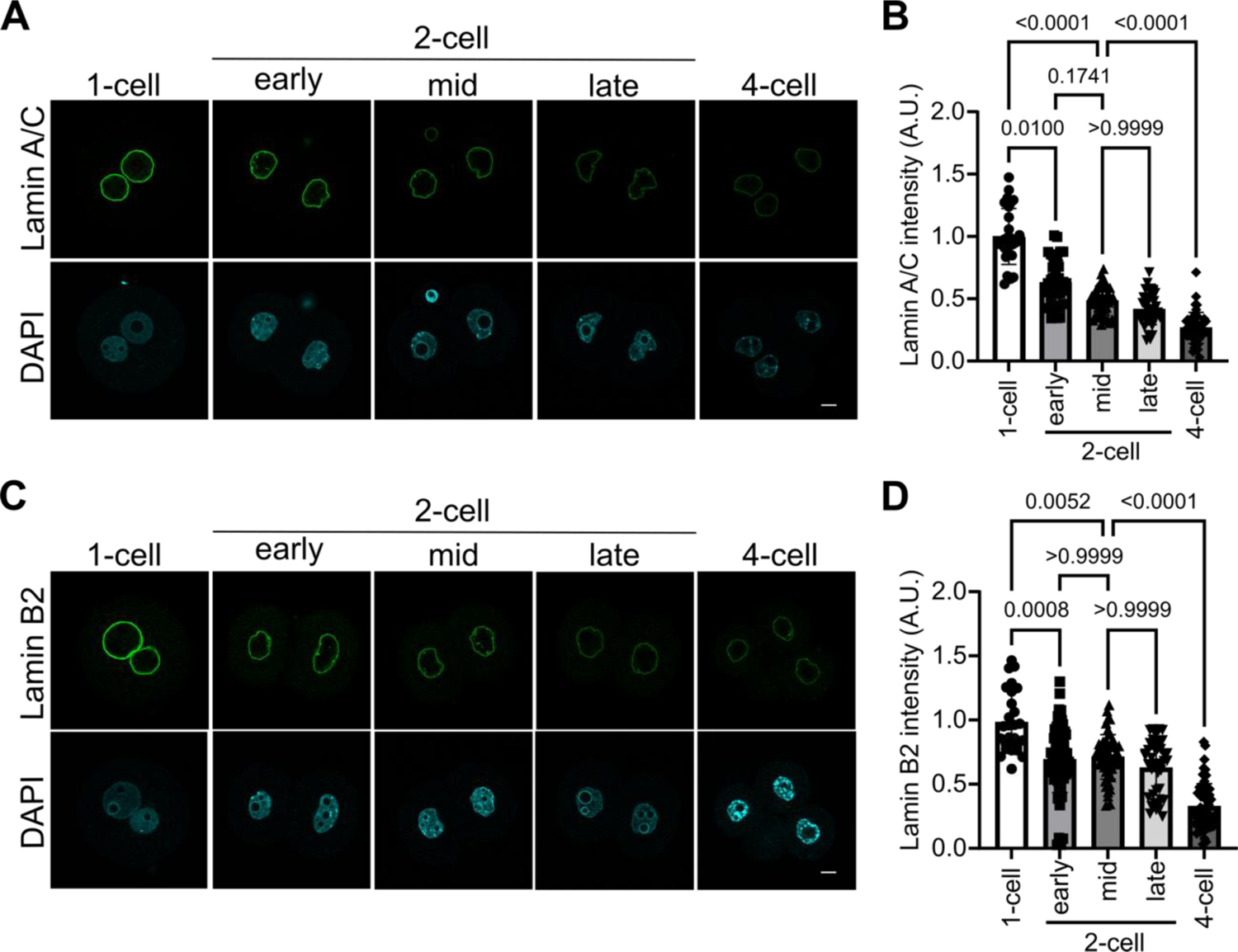
Localization of lamin A/C and lamin B2 at the nuclear membrane of each embryonic stage. (**A, C**) Representative immunofluorescence images of lamin A/C (**A**) and lamin B2 (**C**) (upper panels) and DNA counterstain by DAPI (lower panels) at the indicated developmental stages of mouse embryos. Scale bar, 10 µm (consistent across the images). (**B, D**) Quantification of immuno-fluorescence signals for lamin A/C (**B**) and lamin B2 (**D**) at the nuclear membrane. The integrated signal intensity at the nuclear periphery was scaled to the nuclear circumference length and then normalized to the mean value of the 1-cell stage nuclei (Lamin A/C: *n* = 28, 38, 39, 43 and 44 nuclei, Lamin B2: *n* = 27, 64, 51, 44 and 59 nuclei, respectively). P values were derived from one-way ANOVA and the Kruskal-Wallis test. Mean ± SD are also shown.

**Figure S6.**
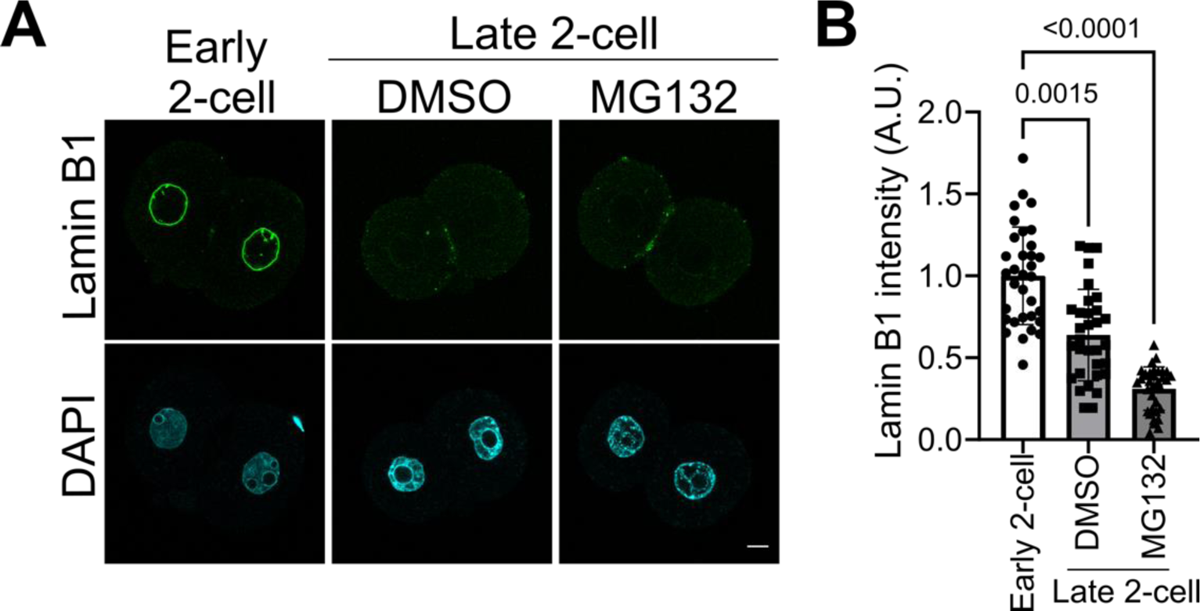
The loss of lamin B1 at the 2-cell stage is not suppressed by inhibition of the proteasome-degradation pathway. (**A**) Representative immunofluorescence images of lamin B1 (upper panels) and DNA counterstain by DAPI (lower panels) in mouse embryos treated with MG132 or DMSO (vehicle control) and fixed at the late 2-cell stage (32 hpi). Immunostaining was also performed for early 2-cell embryos (21 hpi) as a positive control. Scale bar, 10 µm (consistent across the images). (**B**) Quantification of lamin B1 signals at the nuclear membrane. The integrated signal intensity at the nuclear periphery was scaled to the nuclear circumference length and then normalized to the mean value of the early 2-cell stage (*n* = 33, 32 and 35 nuclei). P values were derived from one-way ANOVA and the Kruskal-Wallis test. Bars are Mean ± SD.

**Figure S7.**
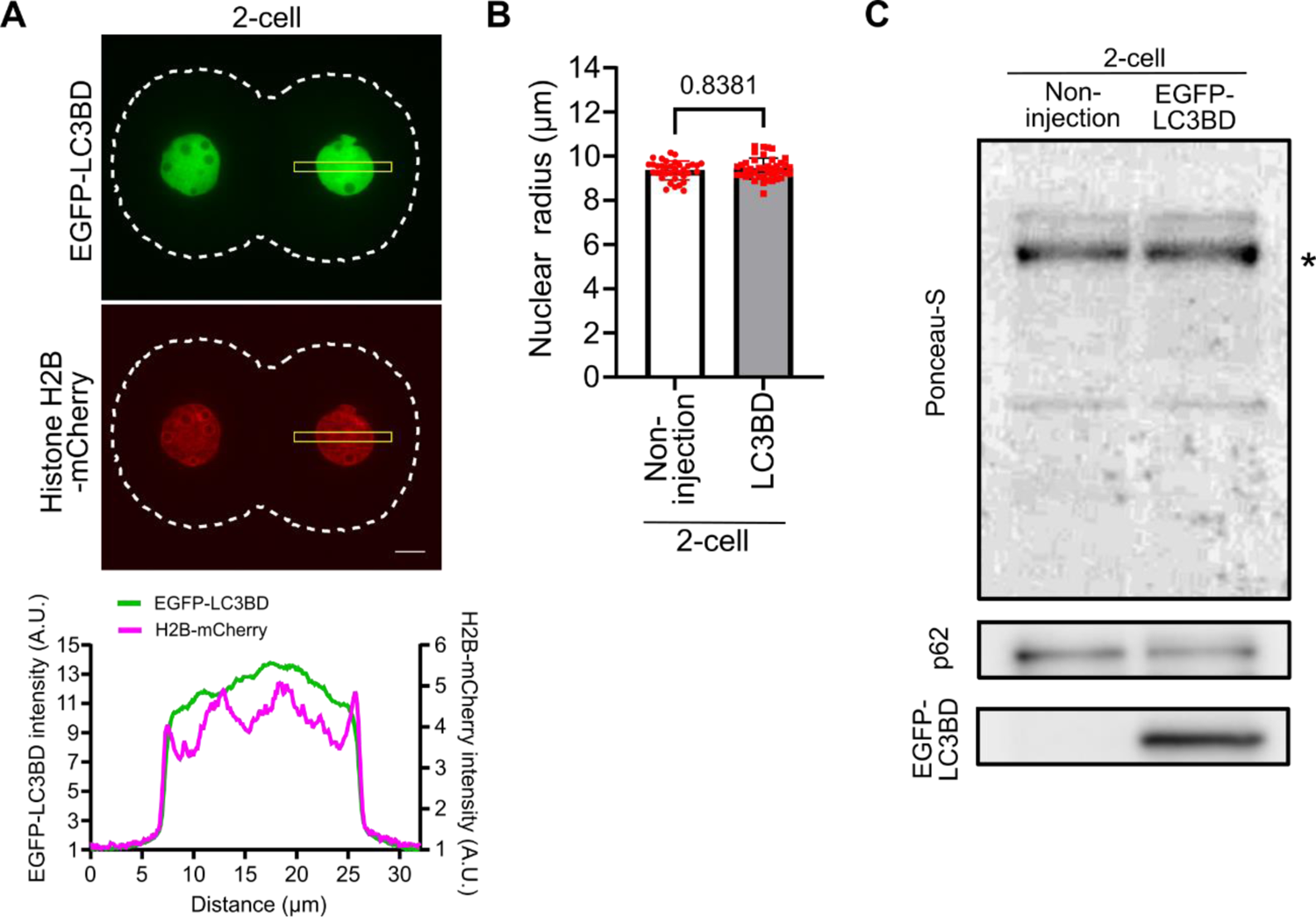
Overexpression of LC3BD does not significantly affect cellular activities. (**A**) A representative confocal live-cell image of a 2-cell stage embryo (24 hpi) expressing the LC3 binding domain of lamin B1 (LC3BD, upper image). EGFP-tagged LC3BD mRNA was injected and expressed in embryos. No detectable accumulation of the LC3-binding domain was observed at the vicinity of the nuclear membrane. H2B-mCherry was co-expressed to visualize nuclear shape (lower image). The bottom profile shows a line-scan generated across the nuclei (yellow highlighted area). White dashed lines indicate cell contour. Scale bar, 10 µm (consistent across the images). (**B**) Comparison of nuclear sizes between control (non-injection) and LC3BD-injected embryos at the 2-cell stage (24 hpi). There was no recognizable difference in nuclear growth between the control and LC3BD embryos (P = 0.83 by Mann–Whitney *U* test). The values in (**B**) are 9.3 ± 0.4 µm (control; *n* = 36 nuclei) and 9.4 ± 0.5 µm (LC3BD; *n* = 42 nuclei) (mean ± SD). (**C**) Immunoblotting results of p62. Lysates of 2-cell stage embryos (32 hpi; 60 embryos per each sample) injected with EGFP-LC3BD mRNA (right lane) and non-injection control (left lane) were blotted. The p62 level was comparable while the LC3-binding domain of lamin B1 was expressed. BSA (asterisk) was mixed with embryos for loading control (Ponceau-S).

**Figure S8.**
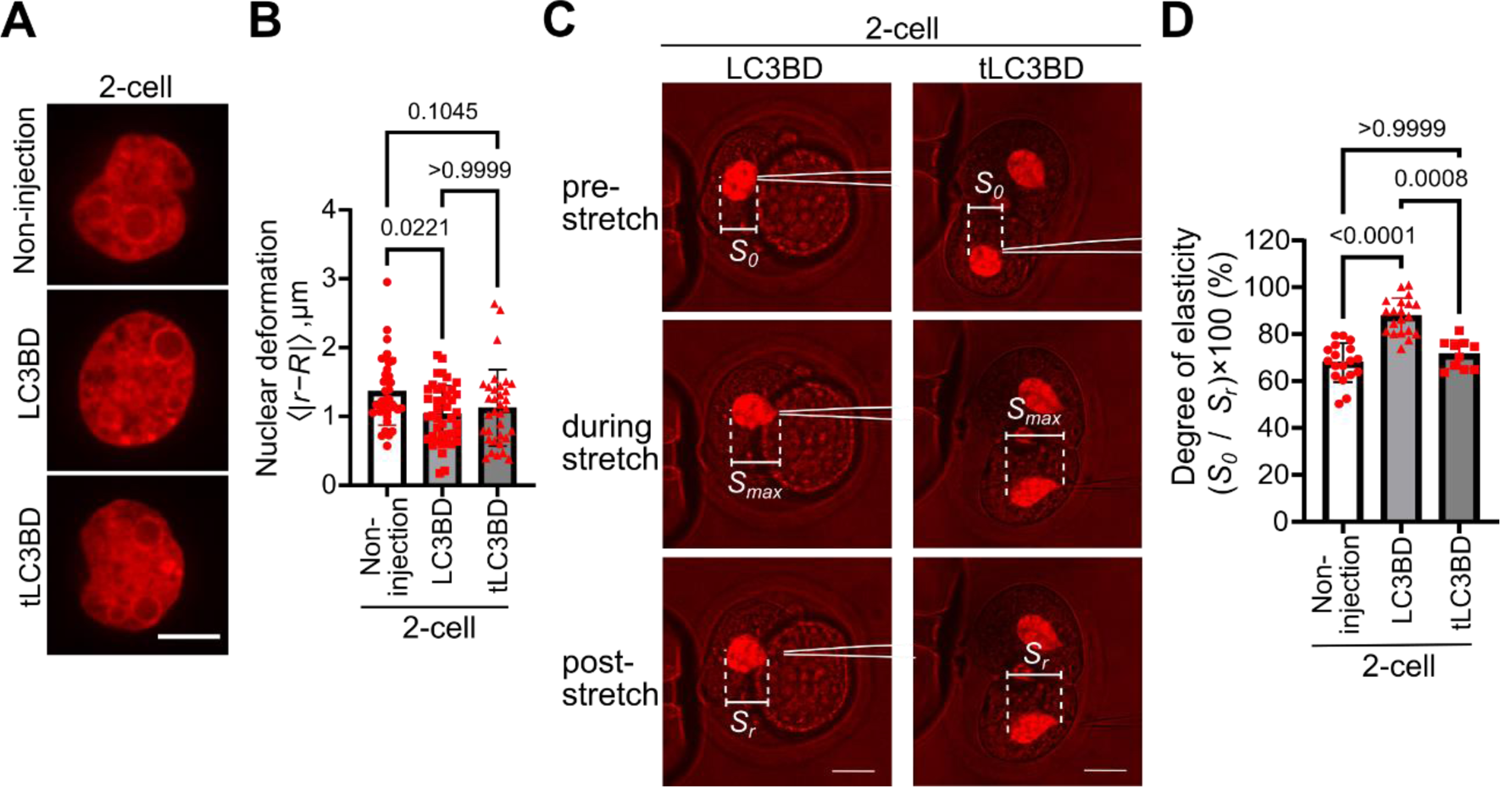
Inhibition of the autophagy-dependent loss of lamin B1 restores the mechanical properties of the 2-cell stage embryonic nucleus. (**A**) Representative confocal live-cell images of nuclei at the 2-cell stage in control embryos (top panel, non-injection) and those injected with LC3BD (middle panel) or tLC3BD (bottom panel; the truncated mutant). H2B-mCherry was used to visualize nuclear shape. Scale bar, 10 µm. (**B**) The magnitude of nuclear deformation was analyzed as in Fig. 1C, yielding the following values: 1.4 ± 0.5 µm (control; *n* = 36 nuclei), 1.0 ± 0.4 µm (LC3BD; *n* = 42 nuclei) and 1.1 ± 0.6 µm (tLC3BD; *n* = 46 nuclei) (mean ± SD). (**C, D**) Nuclear deformability in 2-cell embryos with LC3BD or tLC3BD expression. (**C**) Representative confocal live-cell images of 2-cell stage embryos (24 hpi) injected with H2B-mCherry along with either LC3BD (left panels) or tLC3BD mRNAs (right panels); the magnitude of maximal nuclear deformation was examined using nanocapillaries. Scale bars, 20 µm (consistent across the images). (**D**) The degree of elasticity was determined as in Fig. 1L. The values from each measurement are as follows: 67.8 ± 8.3% (control; reproduced from Fig. 1L), 87.6 ± 7.7% (LC3BD; *n* = 17 nuclei) and 71.3 ± 6.1% (tLC3BD; *n* = 9 nuclei) for degree of elasticity. P values were derived from one-way ANOVA and the Kruskal-Wallis test.

**Figure S9.**
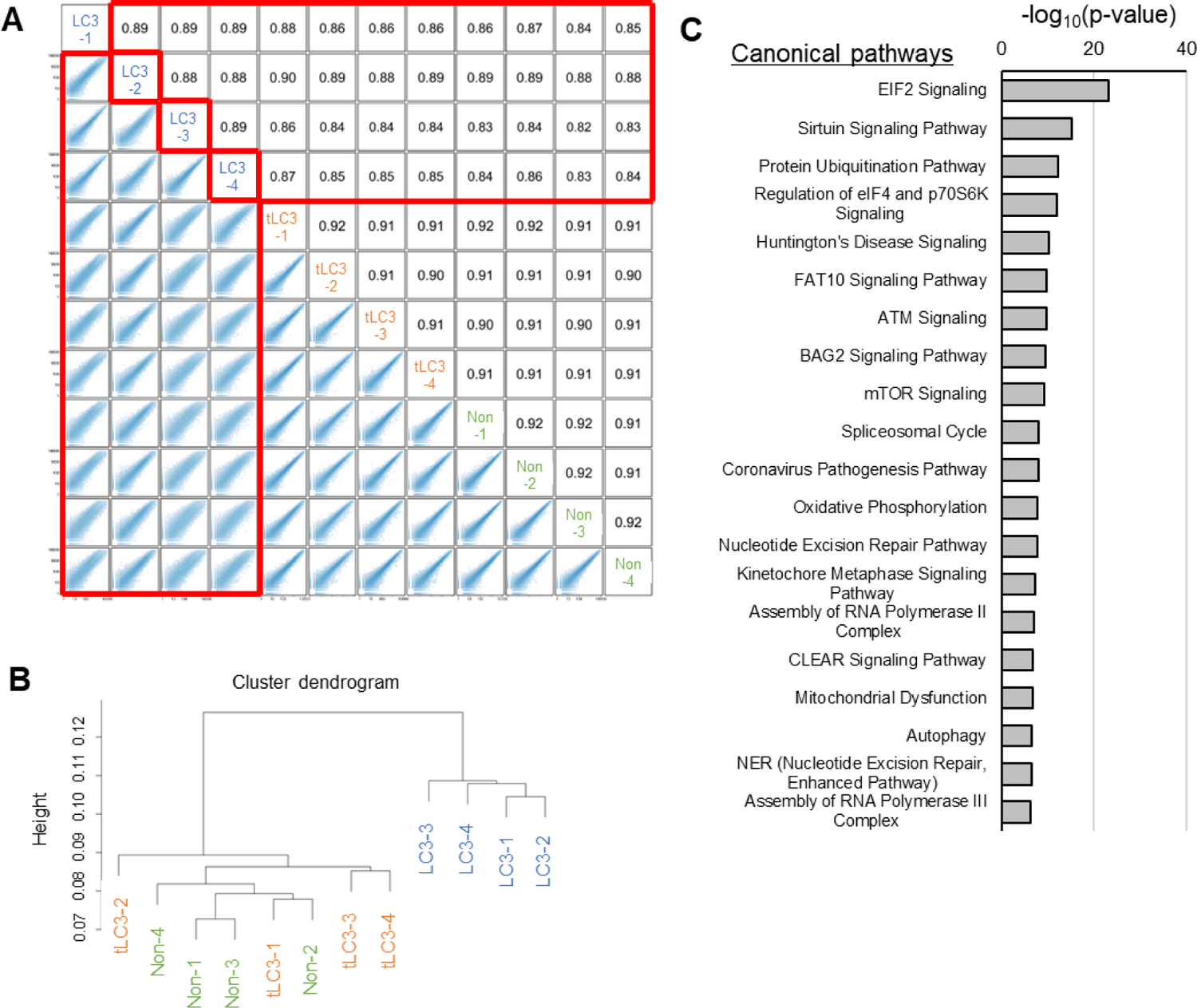
The forced retention of lamin B1 in 2-cell embryos alters transcriptomes. (**A**) Scatterplot and spearman correlation coefficient comparing gene expression profiles among different samples. Biological replicates in each condition are shown such as LC3-1-4 for LC3BD-expressed embryos, tLC3-1-4 for tLC3BD-expressed control embryos, and Non1-4 for non-injection control. Comparisons to LC3 samples are marked by red. (**B**) Hierarchical clustering dendrogram generated using the gene expression profiles of all samples. Spearman correlation was used. (**C**) Canonical pathways predicted by IPA using the list of LC3-downregulated genes (6674 genes in Fig. 3D). Top 20 significant terms are shown (Fisher’s exact test).

**Figure S10.**
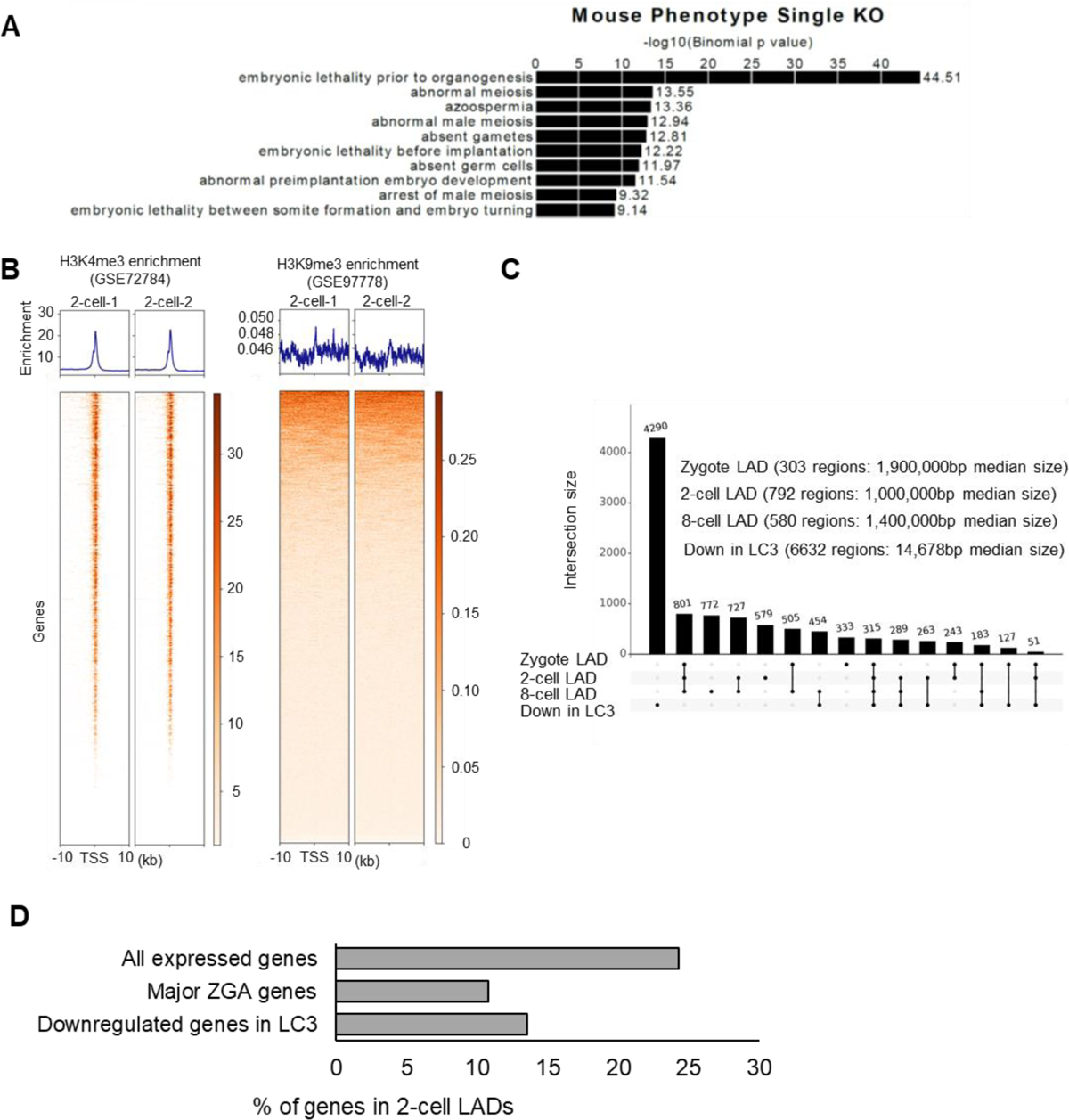
Genes downregulated in LC3BD-expressed embryos are mostly marked with H3K4me3 and not located within LADs in wild type 2-cell embryos. (**A**) Developmentally important genes are enriched in LC3-downregulated genes (4680 genes in Fig. 3D) as revealed by GREAT (Genomic Regions Enrichment of Annotations Tool). (**B**) Heatmaps showing H3K4me3 and H3K9me3 enrichment around transcription start sites (TSSs) of downregulated genes in LC3 embryos (Non vs LC3) in wild type mouse 2-cell embryos. Published ChIP-seq data (*6, 59*) were used to analyze the enrichment. (**C**) The number of genomic regions that overlapped or did not overlap with various regions: zygotic, 2-cell, and 8-cell LADs (*18*) and downregulated genes in LC3 embryos. The number of regions and median size of each genomic feature is shown at top, right. (**D**) Percentages of genes that were associated with 2-cell LADs. The association was regarded when more than 100 bp were overlapped between each gene and 2-cell LADs.

